# Discovery of a Novel Coltivirus in a Newly Identified Bat Bug Species (Heteroptera: Cimicidae) in Cambodia

**DOI:** 10.1101/2025.05.09.652812

**Authors:** Jurre Y. Siegers, Heidi Auerswald, Pierre-Olivier Maquart, Tamara Szentiványi, Julia Guillebaud, Thavry Hoem, Kimhuor Suor, Leakhena Pum, Limmey Khun, Sithun Nuon, Kimlay Chea, Vireak Heang, Kathrina Mae Bienes, Janin Nouhin, Sébastien Boyer, Erik A. Karlsson

## Abstract

Bats and their ectoparasites are significant reservoirs and potential vectors of emerging zoonotic pathogens, yet the viral diversity within bat-associated arthropods remains poorly characterized. This study reports the identification of a novel coltivirus (order *Reovirales*), provisionally designated *Stricticimex coltivirus* (SCCV), in a newly described bat bug species, *Stricticimex phnomsampovensis*, collected from cave-dwelling wrinkle-lipped free-tailed bats (*Mops plicatus*) in Cambodia. Metagenomic sequencing and phylogenetic analysis revealed that SCCV clusters within the *Coltivirus* genus, showing closest similarity to Tai Forest Reovirus (TFRV) previously isolated from African bats. SCCV was detected in 18.4% of examined bat bugs and successfully isolated in VeroE6 cells, with replication confirmed in multiple mammalian cell lines. The discovery of SCCV extends the known diversity and geographic range of *Coltivirus* and highlights bat ectoparasites as overlooked hosts of potentially zoonotic viruses. These findings underscore the importance of integrated One Health surveillance targeting both bats and their ectoparasites to better assess the risk of pathogen spillover in biodiverse regions with high human-animal contact.

**Author Summary:** In this study, we identify a novel Coltivirus, named Stricticimex coltivirus (SCCV), in a newly described bat bug species collected from cave-dwelling bats in Cambodia. Using metagenomic sequencing and phylogenetic analysis, we find that SCCV is closely related to Tai Forest Reovirus, previously identified in African bats. We successfully isolated the virus in mammalian cell lines, suggesting potential to infect vertebrate hosts. This discovery not only expands the known diversity of Coltiviruses, but also underscores the role of bat ectoparasites as underexplored reservoirs of potentially zoonotic viruses. Our findings emphasize the importance of integrated One Health surveillance efforts targeting both bats and their ectoparasites to better assess the risk of virus spillover in regions where human and wildlife habitats overlap.

## Introduction

Bats, as one of the most diverse mammalian groups, have become a key focus in disease surveillance due to their role as reservoirs for pathogens capable of causing human diseases. While ectoparasitic arthropods are well-known vectors of human pathogens in other host groups, their role in pathogen transmission among bats, humans, and other species remains poorly understood. Common bat-associated ectoparasites, including bat bugs (*Cimicidae* and *Polyctenidae*), bat flies (*Nycteribiidae* and *Streblidae*), fleas (*Ischnopsyllidae*), mites (*Acari*), and ticks (*Argasidae* and *Ixodidae*), are hematophagous and occasionally feed on other species, including domestic animals, wild animals, and humans. These ectoparasites are known to carry zoonotic and potentially zoonotic pathogens(1, 2), and therefore they may pose a public and animal health threat(3–8).

In Cambodia, the *Cimicidae* fauna comprises six species **(Table 1**)(9, 10). Among the two species of bed bugs, only *Cimex hemipterus* (Fabricius, 1803) primarily feeds on humans(11–13). The other cimicids reported from Cambodia, including *Aphrania thnotae* (Klein, 1970), *A. vishnou* (Mathur, 1953), *Cimex insuetus* (Ueshima, 1968), *Crassicimex apsarae* (Klein, 1969), and *Stricticimex parvus* (Ueshima, 1968), are specialized bat ectoparasites, like most cimicids species(13). However, two bat bug species, *A. vishnou* (Mathur, 1953) and *S. parvus* (Ueshima, 1968), have been reported to readily feed on humans upon cave entry(9, 14). Given the ability of some of these species to feed on both bats and humans, they serve as potential vectors facilitating pathogen transmission between the two, highlighting the need for further investigation into their public health implications. Additionally, bat guano farming and bat ecotourism increase the frequency of human exposure to bats and their ectoparasites, heightening the risk of pathogen spillover and emerging infectious diseases.

**Table 1:** *Cimicidae* species reported in Cambodia.

| Subfamily | Species | Year of report | Location | Host |
| --- | --- | --- | --- | --- |
| **Cimicinae** | *Cimex insuetus* Ueshima 1968<br>(*Syn. Cimex angkorae* Klein 1969) | 1969(10) | Angkor Wat, Siem Reap | *Tapozous melanopogon bicolor* Temminck(14) |
|  | *Cimex hemipterus* (Fabricius, 1803)<br>(Bed bug, cosmopolitain) | 1969(9) | Pochentong, Chroi Chang War (sic.; Chroy Chang War); Phnom Penh, Kompong Séla (sic.; Kampong Seila); Kampot | Humans |
| **Cacodmiinae** | *Aphrania thnotae* Klein, 1970 | 1970(14) | Ang Sdock, Takeo province, Prek Takong Kandal province | *Scotophilus kuhli gairdneri* Kloss |
|  | *Aphrania vishnou* (Mathur, 1953)<br>(*Syn. Aphrania orientalis* Ferris & Usinger, 1953) | 1969(10) | Prek-Léap, Phnom Penh (Klein, 1969a); Barong region, Phnom Penh; Prek Phnau, Phnom Penh; Phnom Chiso (sic Phnom Chisor) | *Cynopterus sphinx* (Mathur, 1953), *Scotophilus kuhli gairdneri* Kloss (mentioned as *Scotophilus temmincki*), *Cynopterus brachyolus angulatus*; humans(9) |
|  | *Crasscimex apsarae* Klein, 1969 | 1969(9) | Angkor Wat, Siem Reap(9); Vat Nokor (sic. Wat Nokor) Kampong Cham province(14) | *Tapozous melanopogon bicolor* Temminck(9) |
|  | *Stricimex parvus* Ueshima, 1968<br>(*Syn. Strictimex khmerensis* Klein, 1969) | 1969(9) | Vat Nokor in Kampong Cham(14) | *Tapozous melanopogon bicolor* Temminck(9), *Chaerephon plicatus* (Buchannan, 1800); Humans(14) |

Recent advancements in sequencing technologies have significantly increased virus discoveries, particularly in bats and arthropods due to their relevance in potential spillover events to humans and livestock(1, 15–27). One particularly intriguing interface is the bat-arthropod interaction, as these ectoparasites and their microbiome may influence the ecology of infested bats, affecting their health, survival, and behavior(1, 28).

Despite the growing evidence of pathogens in these ectoparasites, surveillance efforts remain biased toward viruses in bats, whereas bacterial pathogens receive more attention in bat-associated ectoparasites(6, 28, 29). Only a few zoonotic viruses have been identified in bat-associated ectoparasites, including Issyk-Kul and Kasokero viruses (Bunyaviridae) in bats and bat ticks(30–33). Additionally, Kaeng Khoi virus, another bunyavirus, was found in *S.x parvus*, *C. insuetus* and free-tailed bats in Thailand, with neutralizing antibodies detected in cave guano miners, suggesting a potential vector role(34, 35). However, beyond these findings, our understanding of the virome of bat-associated ectoparasites, their feeding ecology, vectorial capacity, and public health risks remains limited.

Another group of viruses often detected in ectoparasites and associated with zoonotic infections is the genus of Coltiviruses within the family of *Sedoreoviridae* (order Reoviridae). Currently, there are six member species of Coltiviruses; Colorado tick fever virus (CTFV), Eyach virus (EYAV), Kundal virus (KUNDV), Taï Forest reovirus (TFRV), and Tarumizu virus (TarTV)(36). Coltiviruses are named after its first described member CTFV(37), and are segmented dsRNA viruses, usually transmitted by ticks. CTFV infections in humans are mainly reported in North America and are linked to flu-like symptoms, meningitis and encephalitis(38). The other prominent Coltivirus, EYAV appears to be broadly distributed in Europe with isolations from ticks from Germany(39) and France(40) and associated with human neurological disease through serological evidence(39, 41). Except CTFV and EYAV, all Coltiviruses were identified through molecular surveillance in the last ten years. Most known Coltiviruses or Colti-like viruses were identified and isolated from ticks: *Dermacentor sp.* for CTFV, *Ixodes sp.* for EYAV as well as Shelly headland virus (SHLV) from Australia(16), and Gierle tick virus in Belgium(42), *Hyalomma sp.* for KUNDV from India(43) and Jeddah tick coltivirus from Saudi Arabia(44), and *Haemaphysalis sp.* for TarTV in Japan(45) and O’hara headland virus in Australia(46). Exceptions are Lishui pangolin virus (LSPV) identified in China in 2018 from dead pangolins (*Manis javanica*)(47) and TFRV found 2016 in African free-tailed bats (*Chaereophon aloysiisabaudiae*) in Côte d’Ivoire(48). Out of all Coltiviruses, only CTFV is well characterized including its environmental stability, clinical presentation(49) and transmission through ticks, blood transfusion and vertical transmission from mother to child(50). Available serological diagnostics is limited to in-house ELISAs, IFA and neutralization tests for CTFV(51–55) and EAYV(41, 56, 57).

The discovery of novel viruses at the bat-arthropod interface remains a critical frontier in understanding zoonotic risk, particularly in biodiverse regions where human activities increasingly overlap with wildlife habitats. This study describes the identification of a previously unrecognized *Stricticimex* species in Cambodia and a novel *Coltivirus* in Southeast Asia, expanding both the known diversity of bat ectoparasites and their associated virome. These findings highlight the ecological complexity of bat-ectoparasite-pathogen interactions and underscore the need for targeted surveillance to evaluate their potential role in pathogen transmission and emerging infectious disease threats.

## Materials and Methods

### Collection and processing of bat bugs

The cave of Ta Rumm is located in Sampeu hill, Banan district, Battambang province, north-western Cambodia (13.022 N, 103.095 E). This cave has two vertical entrances, accessible by descending 7-8 meters using a custom-made ladder installed by local guano miners **(Figure 1A & B**). The cave hosts a roost of nearly two million wrinkle-lipped free-tailed bats (*Mops plicatus)* **(Figure 1C**), the sole species documented to inhabit this location(58). Bat guano is harvested in this cave on average twice a month using shovels and a custom pulley system.

**Figure 1:**
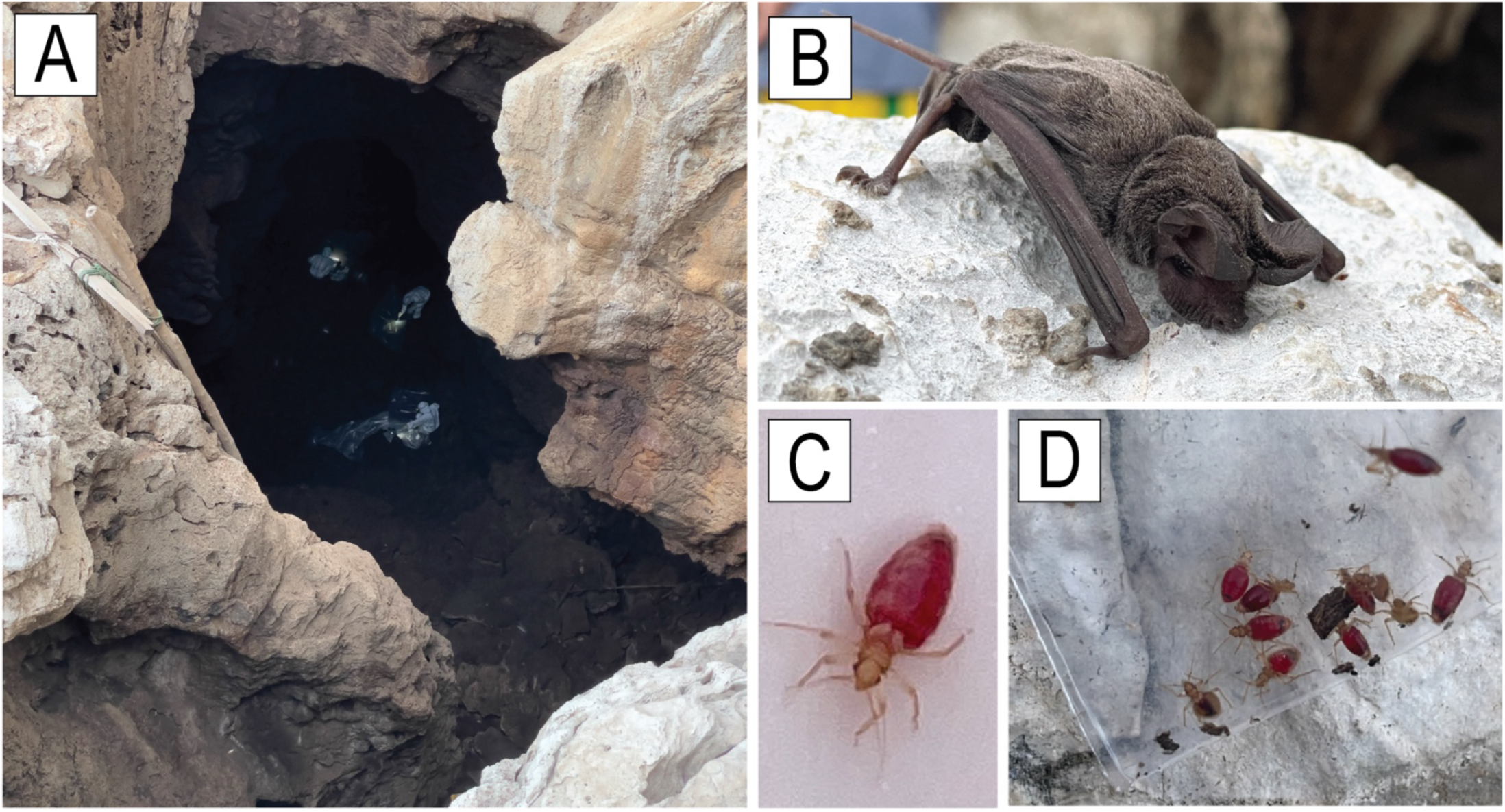
Ta Rumm cave in Phnom Sampov and bat bug collection. (A) Lower entrance of the Ta Rumm cave. IPC field team collecting bat guano and urine inside the cave. (B) Wrinkle-lipped free-tailed bat (*Chaerephon plicatus*). (C-D) blood engorged bat bugs (*Stricticimex phnomsampovensis* Suor & Maquart n.sp). Photos feature only individuals who are authors of this manuscript and were taken with their consent. All images were captured by Jurre Y. Siegers.

In June 2022, the Institut Pasteur du Cambodge (IPC) field team entered the cave through the lower entrance wearing full biosafety level 3-adequate personal protective equipment to collect fresh environmental samples (guano and urine) from bats (**Figure 1A-B**). During this mission, 13 bat bugs (**Figure 1D**) were opportunistically collected, seven were preserved in viral transport medium (VTM) consisting of 2.95% tryptose phosphate broth, 145 mM of NaCl, 5% gelatin, 54 mM Amphotericin B, 106/L U penicillin-streptomycin, 80 mg/L gentamicin (Sigma-Aldrich, Steinheim, Germany), while six bat bugs were stored in 70% ethanol. The bat bugs in VTM were transferred into liquid nitrogen within six hours for transport back to IPC and subsequently stored at −80°C until further analysis. The ethanol preserved bat bugs were kept on ice and handed over to the Medical and Veterinary Entomology Unit at IPC for morphological identification.

The initial identification prompted the need for more specimens. Subsequently, we provided additional tubes to local bat guano collectors for further collection of bat bugs during a harvesting session in April 2023. In total, 80 specimens were stored in VTM, six in 70% ethanol and one was left in no medium for later mounting on a cover slide. These were transported on ice back to IPC within 24 hours and subsequently stored as previously described.

### Morphological Bat Bug Identification

The bat bugs were determined using the determination key provided by Usinger (1966)(59) and the descriptions provided in Ueshima (1968)(60), Klein (1969a, 1969b, 1970)(9, 10, 14). Based on these works, and following examination of closely related species we found that the *Stricticimex* species collected in *Phnom Sampov* is closely related to *S. parvus* and is new to science.

### Speciation (COI, 16s, and 18S) PCR and Sanger sequencing

Parasite species barcoding was performed using three PCR systems targeting invertebrate Cytochrome c oxidase subunit I (COI)(61), 16S and 18S rRNA genes(62–65). Total nucleic acid of individual sample was extracted using Zymo Research Direct-zol RNA MiniPrep kit (Zymo Research, Cat # R2050, CA, USA) according to the manufacturer’s instructions after a homogenization step (MagNA Lyser, Roche). Following reverse transcription using SuperScript III First-Strand Synthesis Super-Mix (Invitrogen, San Diego, CA) according to the manufacturer’s instructions, cDNA was used for the different barcoding PCR protocols.

The COI gene was targeted using primers LC01490-F and HC02198-R. During amplification, the following steps were used: 1 cycle of 94°C for 2 minutes, 40 cycles of 94°C for 30 seconds, 50°C for 30 seconds, and 72°C for 1 minute, and a final extension of 1 cycle of 72°C for 10 minutes. The 16S rRNA gene was targeted with the primers 16S LR-J and 16S LR-N. Amplification was performed with the following conditions: 1 cycle of 94°C for 2 minutes, 35 cycles of 94°C for 30 seconds, 48°C for 40 seconds, and 72°C for 60 seconds, and a final extension of 1 cycle of 72°C for 7 minutes. Finally, the 18S rRNA gene was targeted using pairs of primers 18S-1, 18S-3 and 18S-2, 18S-4. Following conditions were used for both PCR and nested PCR steps: 1 cycle of 94°C for 2 minutes, 40 cycle of 94°C for 30 seconds, 48°C for 60 seconds, and 72°C for 60 seconds, and final extension of 1 cycle of 72°C for 10 minutes.

All amplifications were performed using Invitrogen Platinum™ Taq polymerase and T100 Thermal Cycler, Bio-Rad. All PCR products were subsequently sequenced by Sanger sequencing (Macrogen, Inc., Seoul, Republic of Korea) in both forward and reverse directions using relevant primers from PCR systems (from the nested PCR for 18S amplification).

### Phylogenetic Analysis Cambodian bat bug

The obtained COI, 16S, and 18S sequences were concatenated and aligned with Geneious Prime 2023.2 (Geneious, https://www.geneious.com). Additional sequences of *Cimicidae* species and outgroups were obtained from the National Center for Biotechnology Information GenBank database (**Suppl. Table S1**). A Bayesian consensus tree was created using the MrBayes(66) Geneious plugin, with the General Time Reversible model with gamma distribution and invariant sites(67). Chain length was set to 5,000,000, sampling frequency to 500, and burn-in length to 100,000. The random seed was set to 21,775. Phylogenetic trees were edited using TreeViewer.

### Virus Sequencing

#### Twist Comprehensive Viral Research Panel

The sequencing libraries were generated from homogenized, pooled (n = 4) bat bugs using the combination of Twist Library Preparation Enzymatic Fragmentation (EF) kit 2.0 (#104211) and hybridization with Twist Comprehensive Viral Research Panel (#1035550), according to the manufacturer’s protocol (Twist Total Nucleic Acids Library Preparation EF Kit 2.0 for Viral Pathogen Detection and Characterization Protocol and Twist Target Enrichment Standard Hybridization v1 protocol). Briefly, RNA was extracted using Direct-zol RNA Miniprep Kits (R2053) and the extracted nucleic acid was converted to cDNA using ProtoScript II First Strand cDNA Synthesis kit (E6560S) and Random Primer 6 (S1230S) from New England Biolab (NEB). The NEB Next Ultra II Non-Directional RNA second Strand Synthesis kit (E6111S) was subsequently used to convert the single-stranded cDNA to double stranded DNA (dsDNA). The Illumina TruSeq-compatible libraries were generated using the Twist Library Preparation EF kit and the generated libraries were pooled with 6 libraries per pool. The pooled libraries went to hybridization capture using the Twist Comprehensive Viral Research Panel. Following the enrichment, the enriched libraries were pooled again in equimolar ratios and sequencing with loading concentration of 6pM and spiked with 1% Phix control V3 using Illumina Miseq. The pooled library was diluted and denatured according to the standard Miseq System Denature and Dilute Libraries Guide (Document # 15039740v10), and sequenced to generate paired-end 75bp reads using a 150 cycle Miseq V3 reagent kit (illumina, MS-102-3001). After sequencing, demultiplexed fastq files were generated and analyzed using Genome Detective(68).

#### NovaSeq

Sequencing libraries were generated from Coltivirus infected Vero cell supernatants using the NEBNext® Ultra™ II FS DNA Library Prep Kit for Illumina (New England Biolabs, MA, USA) following the manufacturer’s instructions. First-strand cDNA synthesis was performed using the ProtoScript® II First Strand cDNA Synthesis Kit (New England Biolabs, MA, USA), followed by second-strand synthesis with the NEBNext® Ultra™ II Non-Directional RNA Second Strand Synthesis Module (New England Biolabs, MA, USA). The resulting double-stranded DNA was fragmented to an average size of approximately 400 bp before adapter ligation. Adapter-ligated DNA fragments were then amplified by PCR using the NEBNext® Multiplex Oligos for Illumina. At each step, purification was performed using AMPure XP beads (Beckman Coulter), and DNA quantification was carried out with a Qubit 4.0 fluorometer. Final libraries were sequenced on an Illumina NovaSeq X platform (Macrogen, Inc.), generating paired-end 150 bp (PE150) reads.

### Phylogenetic Analysis Coltivirus

A total number of 96 Reoviridae including species in the genus, *Mycoreovirus*, *Cypovirus*, *Dinovernavirus*, *Oryzavirus*, *Fijivirus*, *Aquareovirus*, *Orthoreoviris*, unclassified Reoviridae, and all species of *Coltiviruses* were obtained from NCBI GenBank (assessed on 2025-01-26). A full list of the GenBank protein accession numbers is provided as Supplementary Table S3. Sequences were aligned with MAFFT v.7.490(69), option L-INS-i and trimmed using trimAL(70), option -gappyout. Phylogenetic trees were constructed using IQ-TREE v.2.0.3(71) using ModelFinder with the best-fit amino acid substitution model Q.pfam+F+I+R6 for RdRp and LG+F+R3 for VP2 chosen according to Bayesian Information Criterion (BIC). Trees were visualized and annotated using FigTree v.1.4.4 (http://tree.bio.ed.ac.uk/software/figtree/) and Adobe Illustrator 2024.

### Real-Time quantitative PCR (RT-qPCR)

A duplex real-time quantitative PCR (RT-qPCR) system was developed to target the viral RNA-dependent RNA polymerase (RdRp) gene. The sequence of this gene was aligned using CLC Genomics Workbench 5.5 with the closest available Coltivirus sequences, - Tai Forest reovirus (accession: KX989543.1) and Kundal virus (accession: NC_055248.1). Based on this alignment, two probes were designed: one targeting the newly identified *Stricticimex Coltivirus* sequence and another targeting the Tai Forest reovirus sequence (**Table 2**). Prior to RNA extraction, samples underwent homogenization using a MagNA Lyser instrument (Roche) with 1.2–1.4 mm ceramic beads (Saint-Gobain, China). Viral RNA was extracted from individual samples using the Direct-zol RNA MiniPrep Kit (Zymo Research, Cat #R2050, CA, USA), following the manufacturer’s instructions. Bat bug RNA samples were then screened using the developed duplex RT-qPCR assay. Each 25 µL reaction mixture contained 5 µL of RNA, 12.5 µL of 2× reaction mix from the SuperScript III One-Step RT-PCR System with Platinum Taq Polymerase (Invitrogen, Darmstadt, Germany), 0.5 µL of a 50 mM MgSO₄ solution (Invitrogen), and 0.3 µL of RNasin ribonuclease inhibitor (20–40 U/µL, Promega). Primers and probes were used at a final concentration of 200 nM. Bat bug RNA samples were screened using developed duplex real-time PCR. A 25µL reaction mix contained 5 µL of cDNA, 12.5 µL of 2x reaction buffer provided with the Superscript III one-step RT-PCR system with Platinum Taq Polymerase (Invitrogen, Darmstadt, Germany) containing 0.5µL of a 50mM MgSO_4_ solution (Invitrogen) and 0.3µL of 20-40U/ µL of RNAsin ribonuclease inhibitor (Promega). Primers and probes were used at final concentrations of 200nM. PCR cycling conditions were as follows: 50°C for 15 minutes followed by 95°C for 5 minutes before 45 cycles of 95°C for 15 seconds, 56°C for 30 seconds and 72°C for 1 minute, using CFX9 Real-Time PCR detection system (Bio-Rad).

**Table 2:**
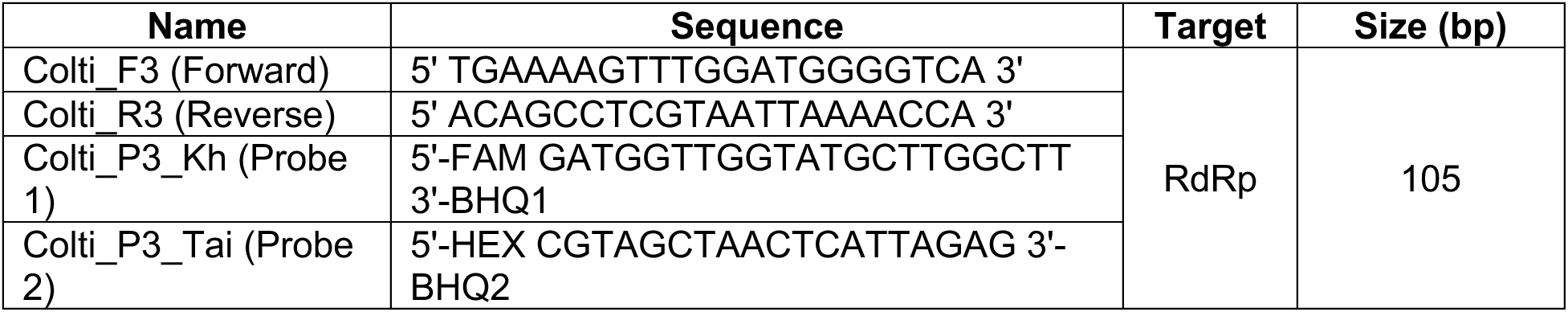
Primers and probe designed to detect Stricticimex coltivirus (SCCV).

### Cell Lines

All cell lines were obtained from either the American Type Culture Collection (ATCC) or the European Collection of Cell Cultures (ECACC), except for the Rhileki cell line, which was provided by G.J. Smith (Duke-NUS, Singapore), and the Aag-2 cell line, provided by J. Pompon (Duke-NUS, Singapore). The Rhileki cell line is a spontaneously immortalized, clonal cell line derived from kidney tissue of a Rhinolophus lepidus bat (NUS-IACUC B01/12), as previously described(72). The invertebrate cell lines Aag-2 (Aedes aegypti) and C6/36 (Aedes albopictus, CRL-1660) were maintained in Leibovitz L-15 medium (Sigma-Aldrich) supplemented with 5% fetal calf serum (FCS; Gibco), 10% tryptose phosphate broth (Sigma-Aldrich), 2 mM L-glutamine (Gibco), and 100 U/mL penicillin-streptomycin (Pen/Strep; Gibco) at 28°C. Vertebrate cell lines were cultured at 37°C with 5% CO₂. Most of these, including BHK-21 (baby hamster kidney, *Mesocricetus auratus*, CCL-10C) and VeroE6 (simian kidney, *Cercopithecus aethiops*, C1008), were grown in Dulbecco’s Modified Eagle Medium (DMEM; Sigma-Aldrich) supplemented with 10% FCS and 100 U/mL Pen/Strep. Caco-2 cells (human adenocarcinoma, HTB-37) were additionally supplemented with 1 mM HEPES (Gibco), while Rhileki cells were supplemented with 1% non-essential amino acids (Gibco) and 1 mM sodium pyruvate (Gibco). HeLa (human adenocarcinoma, CRM-CCL-2) and MDCK (Madin-Darby canine kidney, *Canis familiaris*, CCL-34) cells were maintained in Minimum Essential Medium (MEM; Sigma-Aldrich) supplemented with 10% FCS and 100 U/mL Pen/Strep. For viral infection experiments, the FCS content in the culture medium was reduced to 3% for invertebrate cell lines and 5% for vertebrate cell lines.

### Virus Isolation and Propagation

For virus isolation, homogenized bat bug samples were sterile-filtered using a 0.45 µm filter (Millipore, Burlington, MA, USA) and inoculated to cell monolayers in six-well plates at approximately 80% confluence. Inoculated cells were maintained in DMEM supplemented with 2% FCS, Pen/Strep, and 0.25 μg/mL amphotericin B (Gibco) for seven days. Cell morphology and cytopathic effects were automatically monitored every six hours using an IncuCyte S3 live-cell imaging system (Sartorius, Göttingen, Germany). At the end of the incubation period, cell supernatants were collected by centrifugation and cells were harvested by trypsinization with 0.05% trypsin-EDTA (Gibco). All samples were subsequently analyzed by RT-qPCR. Following initial isolation in VeroE6 cells, the virus was further passaged in this cell line, and virus-containing VeroE6 culture supernatants were used for inoculating growth kinetic experiments.

### Ethics Statement

The field mission was conducted in accordance with Cambodian guidelines and received formal approval from the Ministry of Agriculture, Forestry and Fisheries (MAFF), Ministry of Health National Ethics Committee for Health Research (NECHR #008), local governments, and land/site owners. All necessary permissions were obtained prior to data collection, ensuring compliance with local regulations and ethical standards. The study prioritized biosecurity measures, minimized disruption to the environment and wildlife, and upheld principles of transparency, respect, and responsible data handling.

### Data Availability

Accession numbers for the COI, 16S, and 18S gene sequences of Stricticimex phnomsampovensis are provided in Supplementary Table S1. Sequencing data generated for Stricticimex coltivirus (SCCV) are publicly available in the NCBI Sequence Read Archive (SRA) under BioProject accession number PRJNA1259140.

## Results

### Description of Stricticimex phnomsampovensis

Upon examination, the *Stricticimex* specimens appeared to belong to a new species, closely related to *Stricticimex parvus* and are described below. The updated list of bat bugs in Cambodia, their references, location and known hosts are provided in **Table 1**. The type series is kept in the collection of the Institut Pasteur du Cambodge, Phnom Penh.

### Holotype

**Figure 2**. Male. Head: length: 0.44, slightly longer than wide (0,41), interocular space 5,3 times as wide as eye; Head with 7 setae forming a “y” shape, joining together in the vertex; labrum with 10 pairs of bristles; 5 bristles (3 longer and 2 medium) along inner margin of each eye in addition to about 3 pairs of vertex. Antennae measuring 1.7mm long, always less than 2 mm long; Segments I-IV: 0,15; 0,42; 0,7; 0,43. Second antennal segment as long as width of head, 0,42: 0, 41. Second and fourth segments are subequal. Rostrum 0,48 size of segments I-III: 0,2:0,18:0,1. Pronotum 0,6 wide, twice as wide as long (0,6: 0,28) and straight on the inner sides, narrowed and rounded laterally and posteriorly. Longest bristles at sides as long as those on pronotum. Hemelytral pads are almost twice as large as long (0.37:0.19) and straight on inner sides, narrowed and rounded laterally and posteriorly. Longest bristles at sides about as long as those on the pronotum. Abdomen suboval; second and third segments with 3-4 ill-defined rows of bristles, remaining segments with 2 rows of bristles and with cluster of bristles on terminal segment. Ventral surface with much finer and numerous bristles on the posterior part of each segment. Base of paramere forming an acute angle, then continuing by curving sideways, paramere half as long as the genital segment at base (Fig. 2). Legs long and slender; hind femora about 4,5 times as long as the greatest width, 0.79:0.18; tibiae 2 times longer than femora, 1.6:0.79.

**Figure 2:**
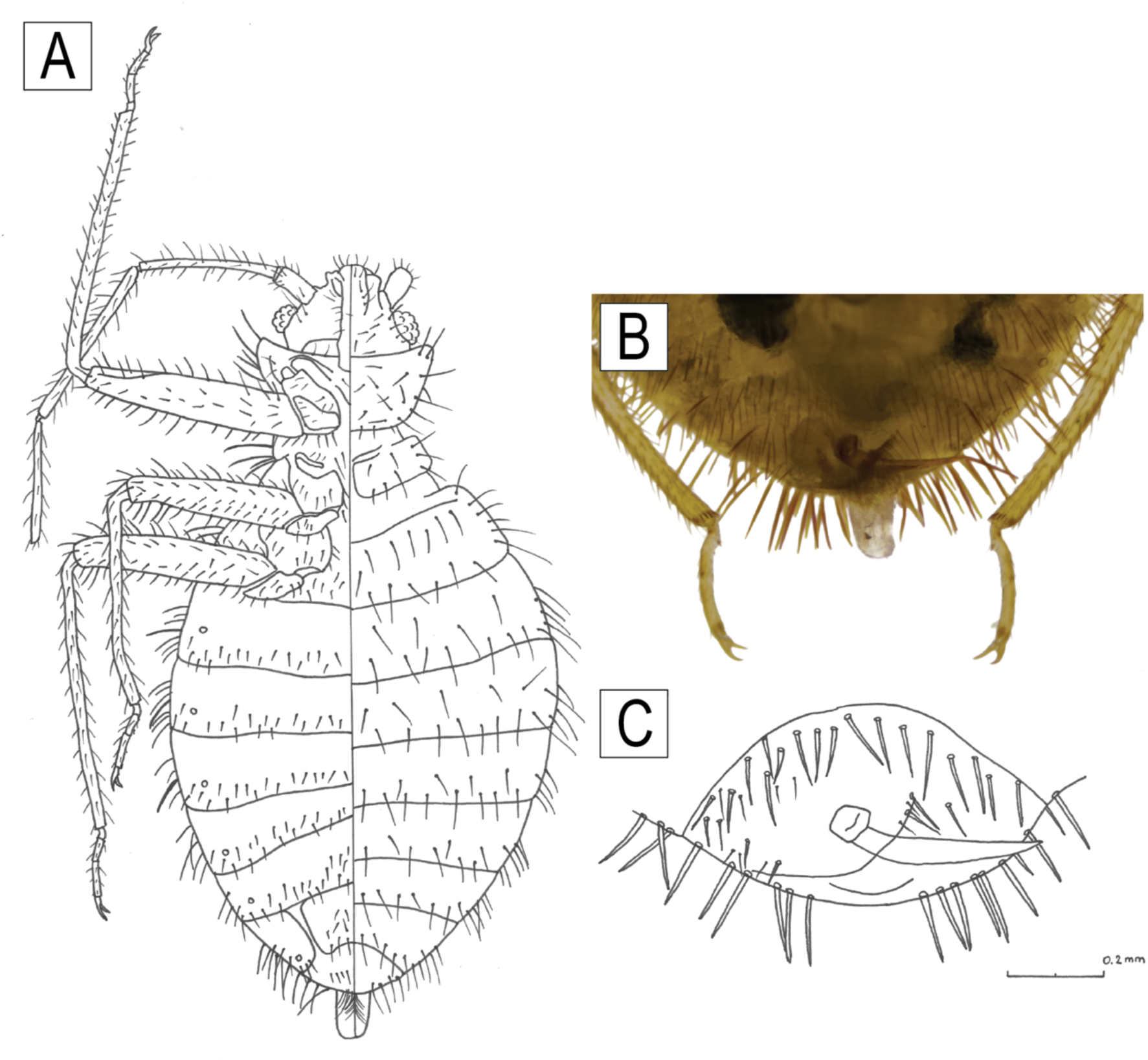
*Stricticimex phnomsampovensis* Suor & Maquart *n.sp* description. (A) Drawing of Stricticimex phnomsampovensis Suor & Maquart n.sp lateral view (left) and dorsal view (right) (Julia Guillebaud, original). (B) Photo Male genital PO Maquart. (C) Male genital drawing (Julia Guillebaud, original).

### Allotype female

Like male. Paragenital sinus sinuate broadly on hind margin of third tergite sublaterally. Ectospermalege broad at opening, directed inwardly and then bent downward. Pronotum width/length: 0,68/0.3. Hemelytras width/length: 0.20/0.38. Hind tibia/femur; 1,86/0,94.

### Etymology

The name refers to the location where this new species was found (Phnom Sampov).

### Ecology

This species was found in a cave dominated by *Mopsplicatus* (Buchanan, 1800), probably feeding off the bats, but they were aggressive, readily feeding on humans entering in the cave (**Figure 1E**).

### Differential diagnosis

The new species, close to *S. parvus* differs by the following characters: (i) Clypeus short and stout while it is more elongated and pointing forward in *S. parvus*; (ii) Hemelytrals pads twice as long as wide, where it is square-shaped in *S. parvus*; (iii) Hind tibiae 2 times longer than femora, while it is 1,5 times longer in *S. parvus*; (iv) Shape of paramere, usually straight or directed downward for *S. parvus* but curving sidewise on the left for *S. phnomsampovensis* with a more acute angle (obtuse angle for *S. parvus)* at the phallobase.

### Authorship

The authors of the new taxon are different from the authors of this article, following the article 50.1 and recommendation 50A of the International Code of Zoological Nomenclature (ICZN, 2012, https://www.iczn.org/the-code/the-code-online/).

### Updated Key to the Species of Stricticimex (revision of Usinger’s 1966 key *in Ueshima*, 1968)

1. Fore femora slightly longer than tibiae. Size small, pronotum 0.6 mm wide. India………. pattoni

(Horvath)

Fore femora is slightly shorter than tibiae. Size medium to large, pronotum 0.7 mm or more in width…. 2

2. Second antennal segment shorter or as long as than width of head……………….. 3

Second antennal segment much longer than width of head………………………….. 6

3. Hind femora less than 4 times as long as wide. South Africa……………… *transversus* Ferris &

Usinger

Hind femora 4 times or more as long as wide 4

4. Second antennal segment as long as the lengh of the pronotum at median line or longer………………….. 5

Second antennal segment shorter than length of pronotum. Interocular space wider than the length of second antennal segment. Size medium. Pronotum 1.0 mm wide. Egypt *namru* Usinger

5. Clypeus pointing forward, hemelytrals pads as long as wide, hind tibiae 1,5 times longer than femora. Obtuse angle at the base of the paramere, continuing almost in a straight line or slightly downward ……………………………………………………………………………………………………………………….. *parvus* Ueshima

Clypeus short and stout, hemelytrals pads twice as long as wide, hind tibiae 2 times longer than femora. Base of paramere forming an acute angle, curving sidewise *phnomsampovensis* **n.sp.**

6. Third antennal segment more than twice as long as fourth. Size large, pronotum 1.0 mm or more in width. South Africa…………………. antennatus Ferris & Usinger

Third antennal segment less than twice as long as fourth. Size smaller, pronotum less than 1 mm wide…………………….. 7

7. Last antennal segment longer than width of head. Longest bristles at sides of pronotum, wing pads and abdomen about 0.37 mm. Kenya…………… intermedius Ferris & Usinger

Last antennal segment subequal to width of head. Longest bristles at sides of pronotum, wing pads and abdomen about 0.31 mm. Democratic Republic of Congo…………… Brevispinosus Usinger

### Phylogenetic analysis of Stricticimex phnomsampovensis

A BLAST search of the COI sequences obtained from bat bugs collected in Ta Rumm Cave revealed the closest match to *Cyanolicimex patagonicus (Haematosiphoninae)*, with a percentage identity of 82.77% and query coverage of 98% (GenBank: MG596833). The 16S sequences showed similarity to *Leptocimex inordinatus (Cacodminae)*, with a 100% percentage identity but low query coverage (65–66%; GenBank: KT592539), indicating potential genetic divergence.

Analysis of the 18S gene fragment revealed high similarity to multiple species across different subfamilies, including *Cimex latipennis* (Cimicinae, 98.9% identity, 100% query coverage; GenBank: KF018720), *Latrocimex spectans* (Latrocimicinae, 98.7% identity, 99–100% query coverage; GenBank: MZ378786), and *Cimex* sp. (Cimicinae, 98.5–98.8% identity, 99% query coverage; GenBank: EU683122). Notably, trimmed sequence comparisons showed 100% similarity among our specimens.

The Bayesian consensus tree, constructed using concatenated sequences of COI, 16S, and 18S rRNA gene fragments, provides strong support for the classification of this newly discovered species within the *Cacodminae* subfamily (**Figure 3**). This phylogenetic placement, combined with the genetic divergence observed, suggests *Stricticimex phnomsampovensis* represents a novel species within bat-associated *Cimicidae* (**Figure 3**).

**Figure 3:**
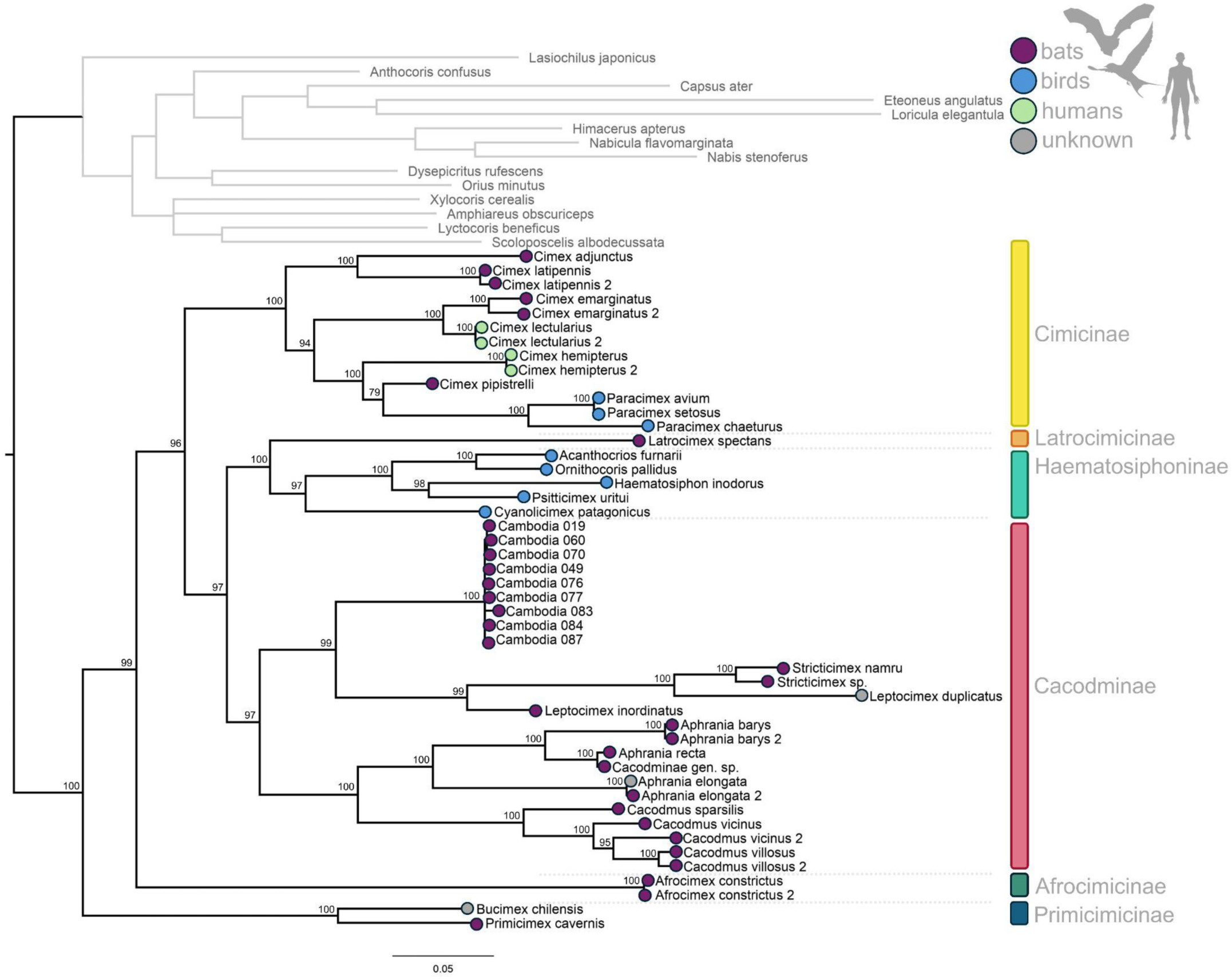
Bayesian consensus tree of family *Cimicidae* (including all six described subfamilies) based on concatenated sequences of COI, 16S, and 18S rRNA gene fragments. GenBank accession numbers are indicated in Supplementary Table S1. The scale bar indicates the number of substitutions per site. Main host groups of each species are indicated with circles (purple: bats; blue: birds; green: humans; gray: unknown host).

### Identification and Phylogenetic analysis of a Novel *Coltivirus* Species in Cambodian Bat Bugs

A novel Coltivirus species, -*Stricticimex coltivirus* (SCCV)-, was identified in *Stricticimex phnomsampovensis* collected from Ta Rumm Cave. The high level of amino acid identity between proteins of SCCV and Tai Forest reovirus (TFRV) and other coltiviruses species suggest that SCCV belongs to the same genus (**Table 4**). Phylogenetic analysis of the available viral genomes revealed that all identified proteins were most closely related to members of the genus *Coltivirus*, which includes the tick-borne pathogenic CTFV, other tick-associated viruses, and TFRV, previously identified in African free tailed bats (**Figure 4**, **Figure 5**). Further in-depth analysis of the RNA-dependent RNA polymerase (RdRp) gene, based on amino acid sequences, showed the highest similarity to TFRV, with nucleotide identity ranging from 74.6% to 75.1% and amino acid similarity between 88.8% and 89.0% (**Figure 4**, **Table 4**). Comparisons of other identified viral proteins revealed amino acid similarities exceeding 70% for VP5, >80% for VP1, VP3, and VP10, and >90% for VP2, VP8, VP9, and VP11, all of which were most similar to TFRV. Notably, VP4 displayed a partially shared similarity with Kundal (**Table 4**). Due to the lack of available nucleotide or protein sequences for VP6, VP7, and VP12 from TFRV, no contigs were generated for these gene segments, leaving their genomic composition unresolved.

**Figure 4.**
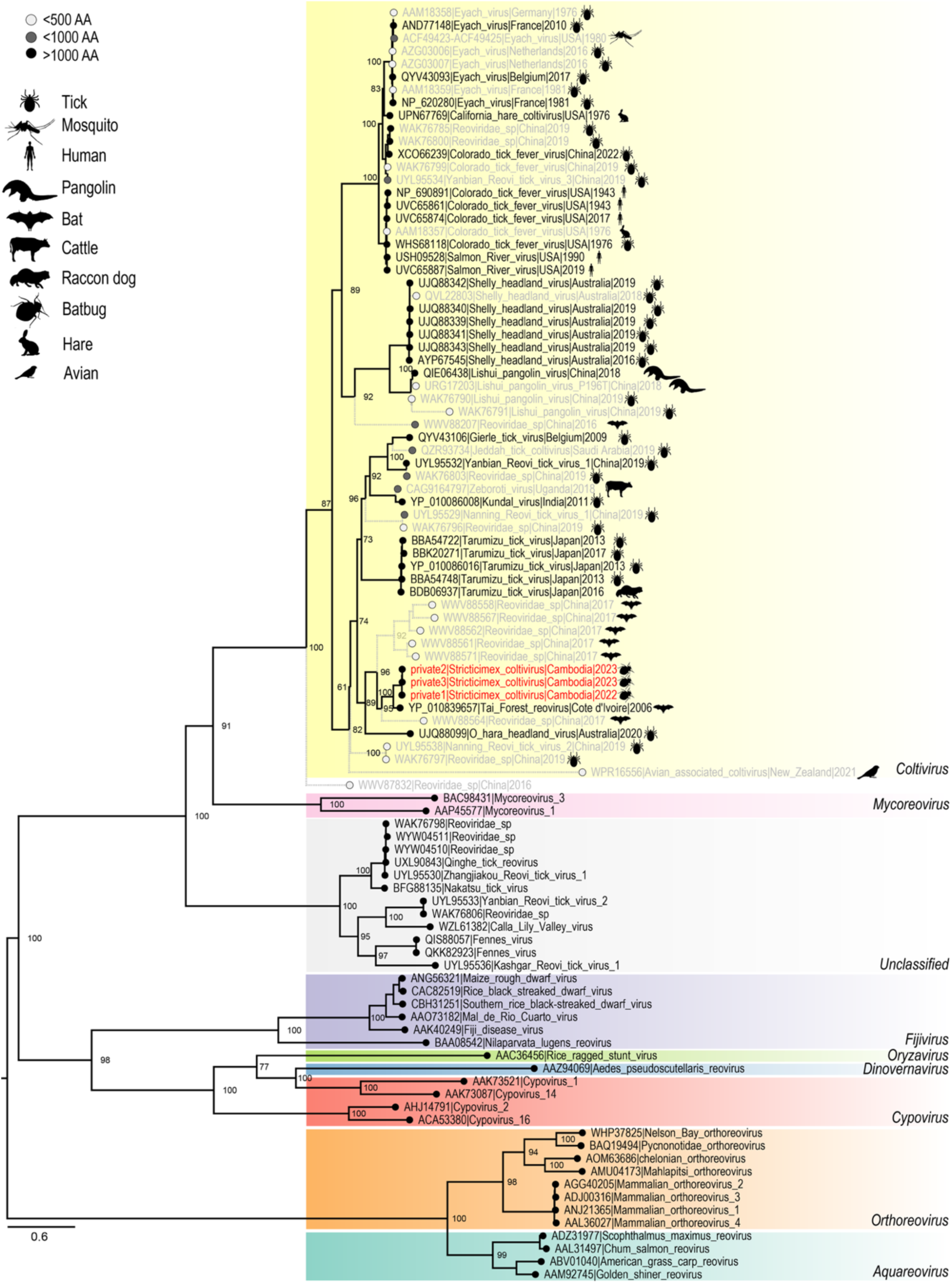
Maximum likelihood tree of the RdRp protein from Coltiviruses and other Reoviridae. Bootstrap values are indicated at key nodes. Stricticimex coltivirus (SCCV) in bold red. Shading associated with the different Reoviridae genera. Tip shading and branch style reflect amino acid lengths of the sequence used to generate phylogenetic tree. Animal symbols within the Coltivirus genus represent the isolation host.

**Figure 5.**
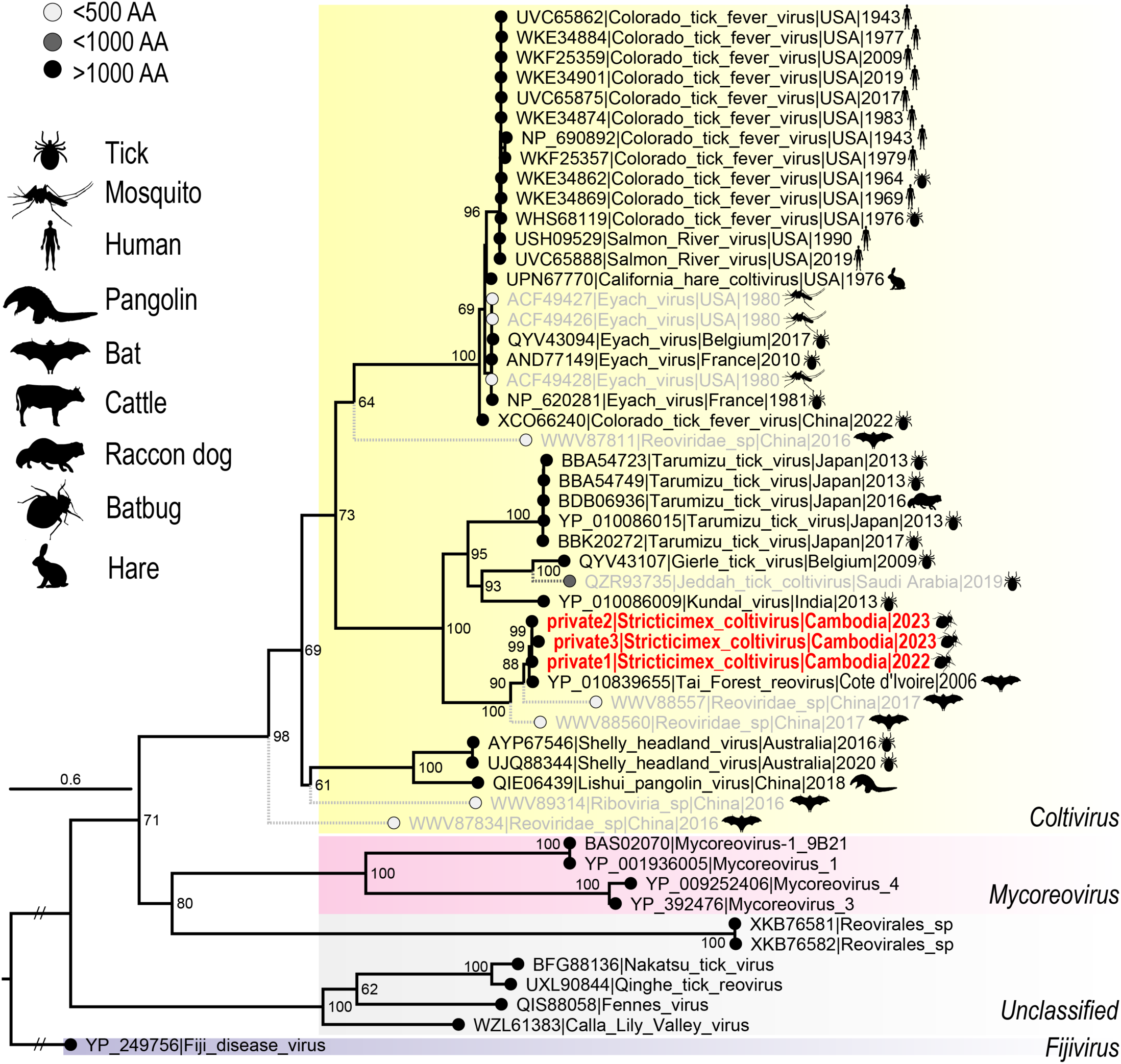
Maximum likelihood tree of the VP2 protein from Coltiviruses and other Reoviridae. Bootstrap values are indicated at key nodes. Stricticimex coltivirus (SCCV) in bold red. Shading associated with the different Reoviridae genera. Tip shading and branch style reflect amino acid lengths of the sequence used to generate phylogenetic tree. Animal symbols within the Coltivirus genus represent the isolation host.

**Table 3:**
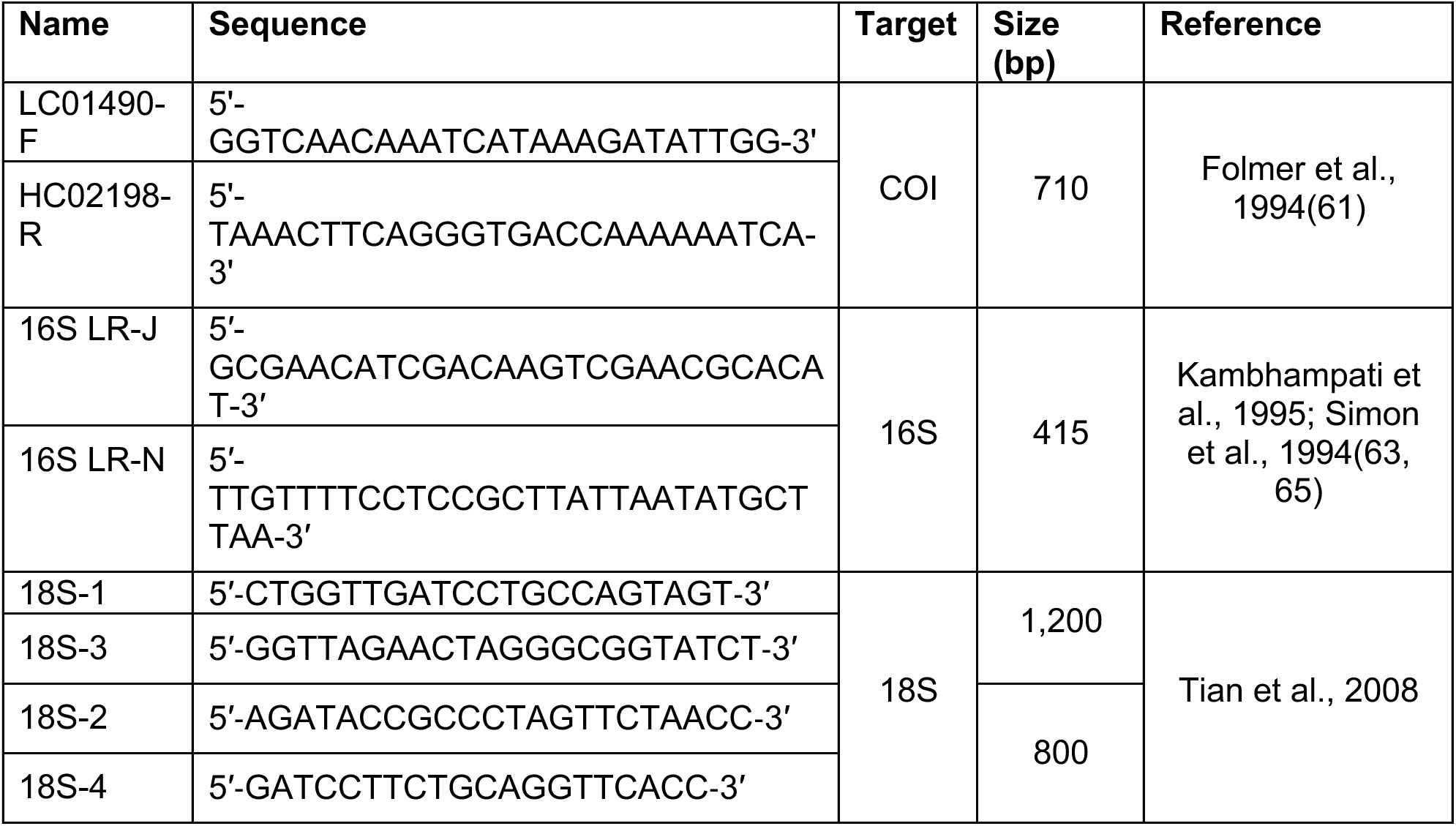
Primers used for species identification.

**Table 4.**
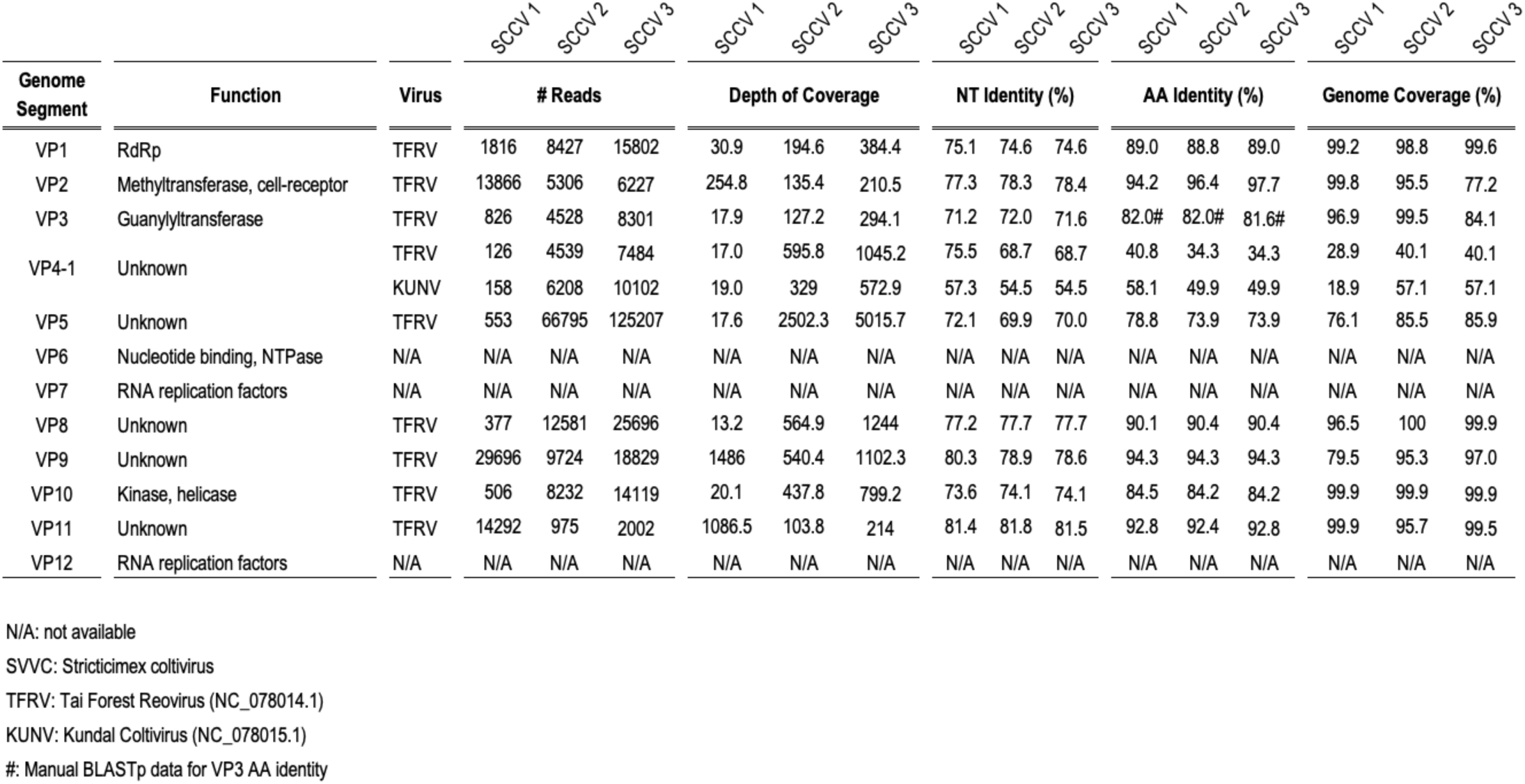
Nucleotide and amino acid similarities between Stricticimex coltivirus, Tai Forest Reovirus and Kundal virus.

Phylogenetic analyses of the conserved RdRp and VP2 protein sequences revealed that the SCCV clusters within a distinct clade alongside known *Coltivirus* and *Coltivirus*-like viruses detected in bats (**Figure 4**, **Figure 5**). Notably, SCCV is most closely related to *Tai Forest reovirus* (TFRV), originally isolated in 2006 from free-tailed bats (*Mops aloysiisabaudiae*) in Côte d’Ivoire. More distantly, and with greater uncertainty due to partial RdRp and VP2 sequences, SCCV also shares similarities with *Coltivirus*-like viruses detected in *Taphozous melanopogon* bats detected in Guangxi, China in 2017.

The strong bootstrap support for the clustering of SCCV sequences with TFRV indicates a relatively close evolutionary relationship (**Figure 4**, **Figure 5**). Phylogenetic analysis of the RdRp and VP2 proteins further reveals that bat-associated Coltiviruses form a monophyletic clade within the Coltivirus genus. However, the presence of partial protein sequences introduces some uncertainty at certain branching points and may not fully capture the true evolutionary relationships.

Taken together, the sequence data, gene segment identities, and phylogenetic placement strongly support SCCV as a novel *Coltivirus* species, expanding our understanding of bat-associated reoviruses. Further studies are needed to fill phylogenetic gaps and refine its evolutionary history and clarify its relationship to other *Coltivirus* members and whether bats and bat-associated ectoparasites occupy a phylogenetic niche within the *Coltivirus* genus.

### Coltivirus prevalence in Cambodian bat bugs

A total of 87 individual bat bugs collected from Ta Rumm Cave in April 2023 were screened for Coltivirus using a newly developed real-time PCR assay targeting the RdRp gene. Sixteen (18.4%) tested positive, with Ct values ranging from 23.02 to 40.58 (mean Ct: 30.18). All positive samples were identified as the SCCV based on a specifically designed probe, while no samples tested positive for Tai Forest reovirus (TFRV).

### Virus Isolation

Virus isolation was attempted using homogenized bat bugs to inoculate C6/36 (*Aedes albopictus*), BHK-21 (*Mesocricetus auratus*), Rhileki (*Rhinolophus lepidus*), and VeroE6 (*Chlorocebus sabaeus*) cell lines. Initial inoculations included samples with Ct values ranging from 23.02 to 40.58. Successful virus isolation was achieved in VeroE6 cells from samples with Ct values of 23.02 and 25.29. The presence of the virus was confirmed in the culture supernatant across three subsequent passages using the SCCV specific RT-PCR assay.

### Virus Growth Kinetics

The replication dynamics of one SCCV isolate was assessed in invertebrate (Aag-2, C6/36) and vertebrate (BHK-21, Caco-2, MDCK, Rhileki, VeroE6) cell lines over 12 days. Culture supernatants and cells were analyzed by RT-PCR to detect the presence of SCCV (**Figure 6**). Viral entry was observed in all tested cell lines at some point during the experiment (**Figure 6B**). However, productive viral replication, indicated by the release of virus into the culture supernatant, was only detected in Caco-2, HeLa, BHK-21, and VeroE6 cells. Among these, BHK-21 and VeroE6 cells exhibited the highest viral titers, with replication increasing over time (**Figure 6A**).

**Figure 6.**
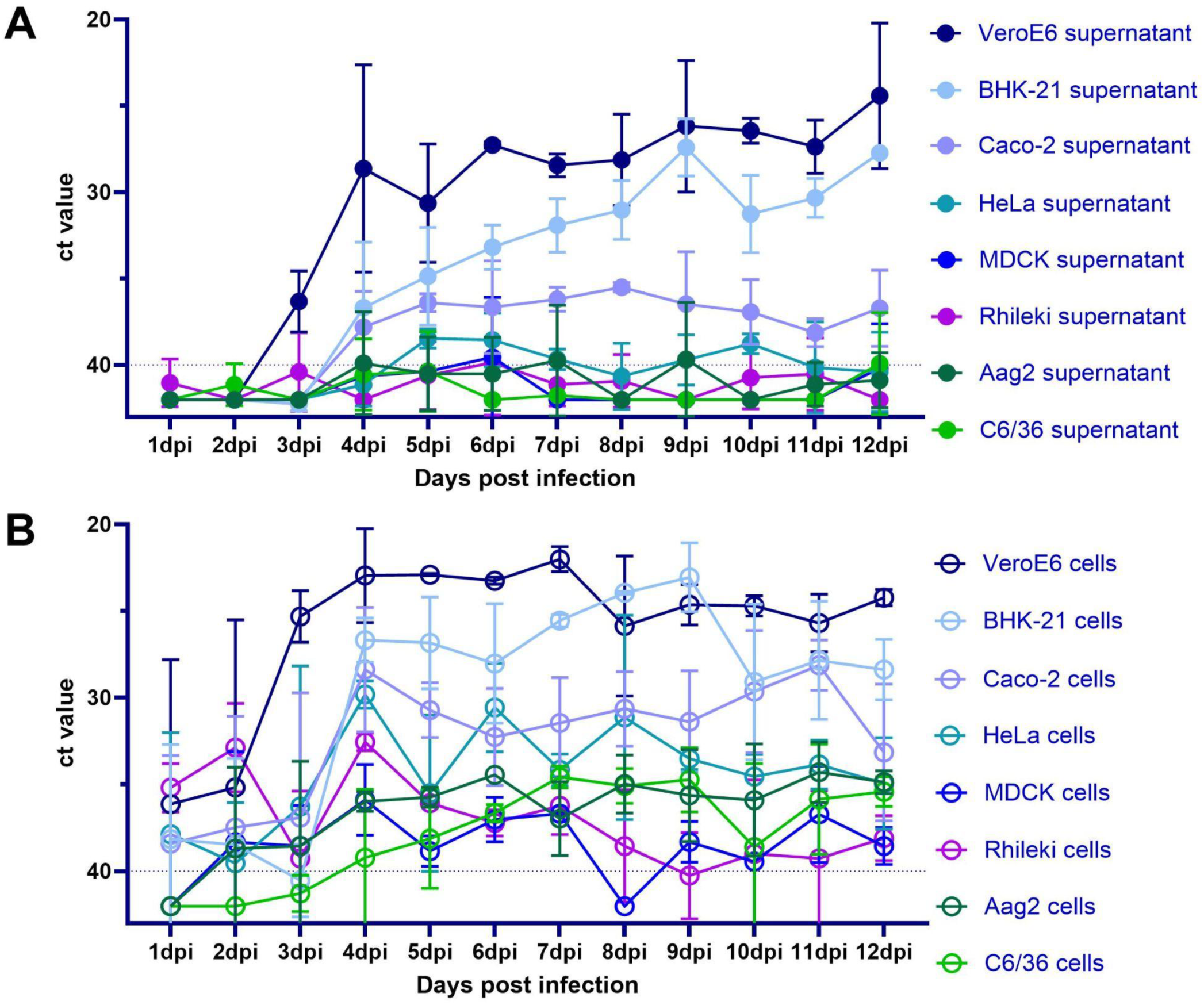
In vivo growth kinetics of Stricticimex coltivirus (SCCV). Various invertebrate (shades of green) and vertebrate (shades of blue) cell lines, including a bat cell line (in violet), were inoculated with the SCCV isolate (passage 3 in VeroE6) for 1 hour at 37°C. Supernatants (A) and cells (B) were collected every 24 hours and analyzed by RT-PCR. Experiments were conducted in biological duplicates. The Ct values are plotted inversely to intuitively demonstrate viral growth.

## Discussion

We report on the identification of a novel coltivirus species in a previously unknown bat ectoparasite, - *stricticimex*-, species in Cambodia highlighting the importance of understanding potentially zoonotic viruses associated with bat ectoparasites. In recent years, several Coltiviruses have been identified in Asia(73), however, to the best of our knowledge, the Coltivirus described in this study marks the first of its genus to be identified in Southeast Asia. Given the fact that some members of the coltivirus genus can cause human disease highlights the importance of understanding potentially zoonotic viruses associated with bat ectoparasites. Phylogenetic analyses suggest that the Stricticimex Coltivirus and Tai Forest Reovirus (TFRV) may share a recent common ancestor, showing limited evolutionary divergence at the protein level despite significant geographic separation. TFRV was isolated from the blood of African free-tailed bats (*Chaereophon aloysiisabaudiae*) in Côte d’Ivoire in 2006 and capable of causing cytopathic effects on C6/36 insect cells and on various mammalian cell lines (VeroE6, human cell lines MRC-5 and Hep2, and a fruit bat cell line originating from *Rousettus aegyptiacus* (R05T))(48). Although both TFRV and SCCV were able to infect and replicate in several mammalian, -including human-, cells, it remains to be determined whether both viruses are able to infect, replicate and cause disease in humans.

The Stricticimex Coltivirus was identified in bat bugs present at a large colony of cave dwelling wrinkle lipped free-tailed bats. Bat parasites are often considered to be host specific due to the ecological isolation of bats or the associated life history strategies of these parasites(74) highlighting the possibility of a similar virus-host dynamic as for TFRV. TFRV is the only Coltivirus reported from bats, with no detection in ticks so far, suggesting a potentially unique virus-host relationship. The prevalence of the SCCV in the collected bat bugs from Ta Rumm cave was notably high at nearly 20%. Given that this study was limited to two sampling events within one year, more extensive surveillance is needed to determine if similar prevalence rates are observed in other caves and during different seasons, and to understand if these rates are influenced by factors such as the bat birthing period or seasonal variations in ectoparasite infestations, similar to patterns observed with coronaviruses(75) and bat flies in other studies(76). As SCCV was identified only in bat bugs, it remains to be investigated if this is primarily a bat or a bat bug virus and whether it has the potential to infect other cave-dwelling species such as rodents. In addition, Ta Rumm cave as well as other caves in the area are used for guano harvesting increasing the chance of zoonotic pathogen transmission to humans either directly from bats or from bat ectoparasites. Future serological studies should elucidate whether guano farming practices increase the risk of coltivirus exposure. In addition, bat ecotourism, such as observation of bat mega colonies exodus at dusk, visits to pagodas inside bat caves, can increase the risk of zoonotic pathogen exposure and transmission by bringing humans into close proximity with bat colonies, thereby facilitating opportunities for direct or indirect contact with bats and bat excreta that may harbor zoonotic pathogens. The induction of CPE in VeroE6 cells and SCCV’s close phylogenetic relationship to human pathogenic viruses suggest its potential to infect humans. Febrile illnesses, like those caused by CTFV, are common in tropical regions and often go undiagnosed. In remote areas in Cambodia, where guano collectors enter bat-inhabited caves, limited healthcare access and a lack of diagnostic testing could mean that an SCCV-like disease remains undetected. Together, this necessitates further research to explore the virus’s reservoir, host range, cross-species transmission capabilities and serological studies into human exposure.

Merely identifying these viral sequences is not sufficient for comprehensive risk assessments. Detailed *in vitro* and *in vivo* studies are required to evaluate their cross-species transmissibility, prior exposure in humans and other animals, and pathogenicity. Such efforts should adopt a One Health approach, integrating human, animal, and environmental health to achieve a holistic understanding of virus ecology. A proactive surveillance, rather than waiting for outbreaks to occur, can potentially detect and prevent future zoonotic spillovers, thereby safeguarding global health. Additionally, this work highlights the importance of traditional and molecular taxonomy of hematophagous parasites, which could further enhance our understanding of potential vectors for animal and human pathogens. Particularly in tropical regions, there are still a number of bat-associated parasites likely undescribed, which could serve as vectors for both known and unknown animal and human-associated diseases. Future studies should pay particular attention to those closely related to species that occasionally feed on humans or other non-chiropteran hosts, such as *Leptocimex* species. These behavioral traits may indicate a broader host range in related species, including *Sticticimex* spp., suggesting their potential role in interspecies pathogen transmission.

The bat bugs analyzed in this study belong to the Cacodminae subfamily, with *S. phnomsampovensis* exhibiting a close phylogenetic relationship to other *Stricticimex* and *Leptocimex* species, as supported by both molecular and morphological data. These phylogenetic findings align with previous studies that clarify relationships among Cimicoidea subfamilies(13, 64, 77). Species within Cacodminae, including *Leptocimex* and *Stricticimex*, are obligate blood-feeding ectoparasites primarily associated with bats from the Molossidae and Vespertilionidae families across Asia, Africa, and Europe(78). The species of Cacodminae subfamily, including species such as *Leptocimex* and *Stricticimex*, are blood-feeding ectoparasites predominantly targeting bats of the Molossidae and Vespertilionidae families across various regions including Asia, Africa and Europe(78). Given the broader context, where tens of thousands of new arthropod species are described annually(79), approximately 14,000 species of arthropods across over 400 genera are known to have developed the capacity to feed on vertebrate blood, including specific groups like certain mosquitoes, ticks, fleas, and some flies, such as sandflies and horseflies. This represents a small fraction of the total arthropod species, indicating that blood-feeding (hematophagy), is a relatively rare trait among arthropods(80). However, the arthropods known for hematophagy have been intensely studied due to their importance in transmitting pathogens.

The discovery of a novel bat bug species and its associated virus underscores the critical need for One Health surveillance, emphasizing the interconnectedness of ecosystems and the potential for zoonotic disease emergence. This study demonstrates that with field access, modern analytical methods, and cross-disciplinary collaboration, pathogen detection in high-risk areas can proactively identify emergence risks. By fostering a comprehensive understanding of pathogen presence, emergence, and spread, One Health surveillance supports proactive public health strategies, mitigating outbreak risks and enhancing global health security(81).

## Author Contribution

The study was conceptualized by H.A., P.-O.M., J.Y.S., J.G., T.H., J.N., S.B., and E.A.K. Data curation and formal analysis were carried out by H.A., P.-O.M., T.S., J.Y.S., J.G., T.H., K.S., L.P., L.K., S.N., K.C., V.H., K.M.B., and J.N. Funding acquisition was secured by J.N., S.B., and E.A.K. The investigation involved H.A., P.-O.M., J.Y.S., J.G., T.H., K.S., L.P., L.K., S.N., K.C., V.H., K.M.B., J.N., S.B., and E.A.K., while methodology development was undertaken by H.A., P.-O.M., T.S., J.Y.S., J.G., T.H., V.H., K.M.B., J.N., S.B., and E.A.K. Project administration was managed by H.A., P.-O.M., J.Y.S., J.N., S.B., and E.A.K., and resources were provided by H.A., P.-O.M., T.S., J.Y.S., J.G., T.H., J.N., S.B., and E.A.K. Supervision was conducted by H.A., P.-O.M., J.Y.S., J.G., T.H., S.B., and E.A.K., with validation and visualization supported by H.A., P.-O.M., T.S., J.Y.S., J.G., T.H., V.H., J.N., S.B., and E.A.K. The original draft was written by H.A., P.-O.M., T.S., J.Y.S., J.G., T.H., J.N., S.B., and E.A.K., and all authors contributed to the review and editing of the manuscript.

## Acknowledgements

We gratefully acknowledge the dedicated field teams of Institut Pasteur du Cambodge for their invaluable efforts in sample collection and logistical support. We also thank the Institut Pasteur du Cambodge Virology Unit team for their technical assistance and laboratory expertise. Special appreciation is extended to the guano farmers at the bat caves, whose cooperation and support made this study possible. We ly acknowledge all relevant authorities from the Ministry of Agriculture, Forestry, and Fisheries and the Ministry of Environment for their great support in facilitating this work. We are especially thankful to the Department of Wildlife and Biodiversity under the Forestry Administration for their continuous assistance and collaboration on ongoing studies in wildlife in Cambodia, which was essential for the success of this study. Special appreciation is extended to the guano farmers at the bat caves, whose cooperation and support made this study possible.

## Supplementary Material

**Supplementary Table S1:**
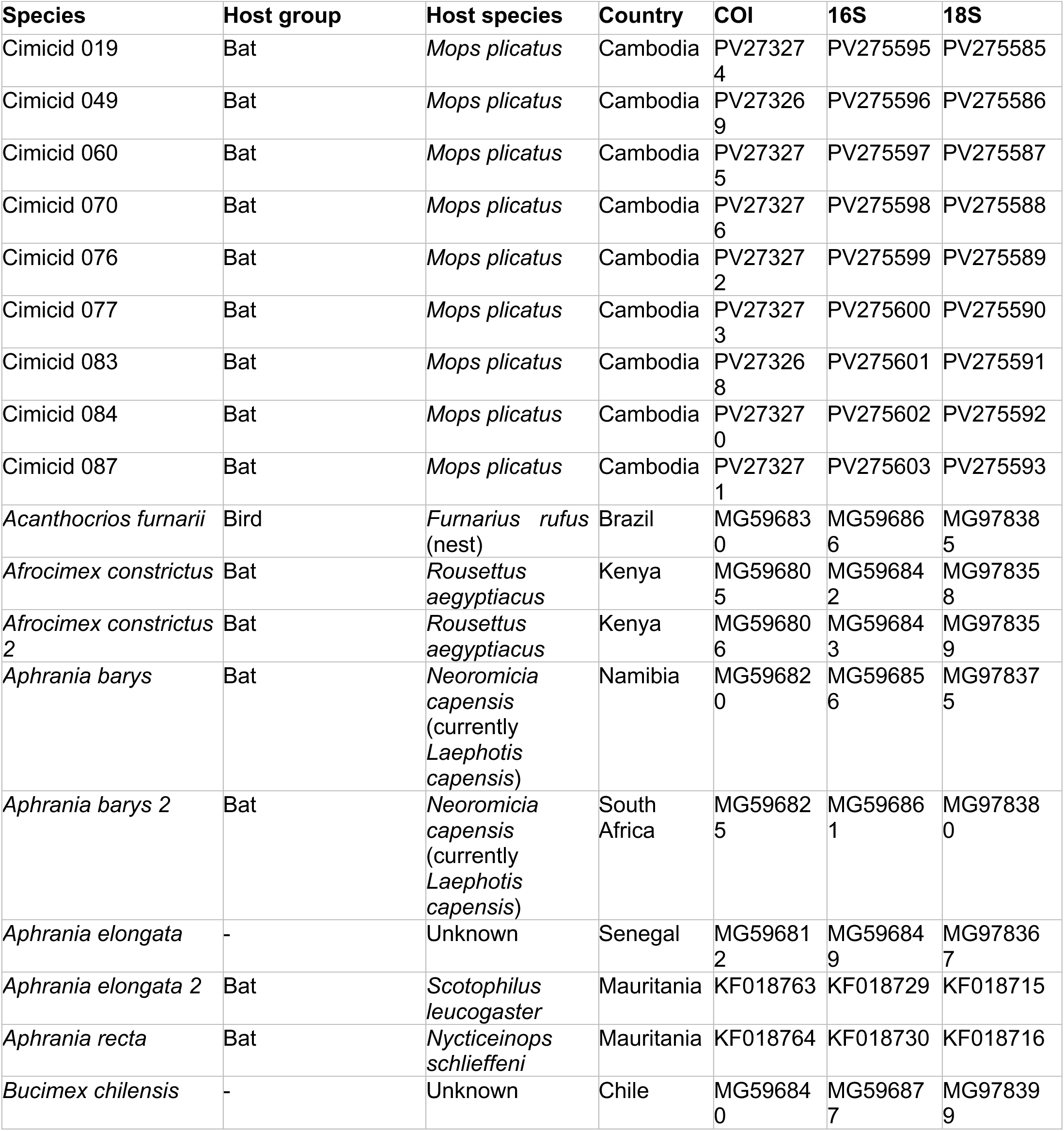
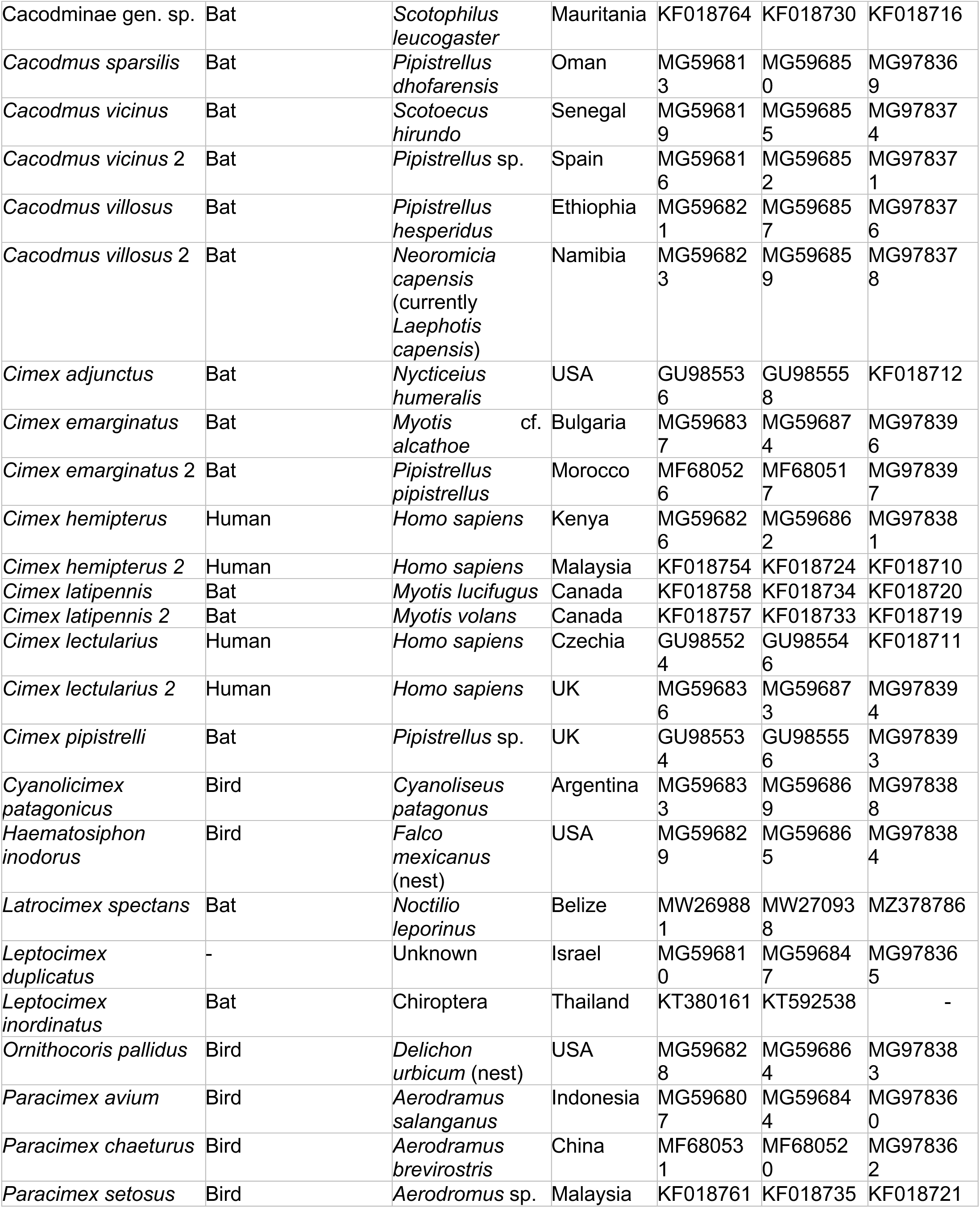
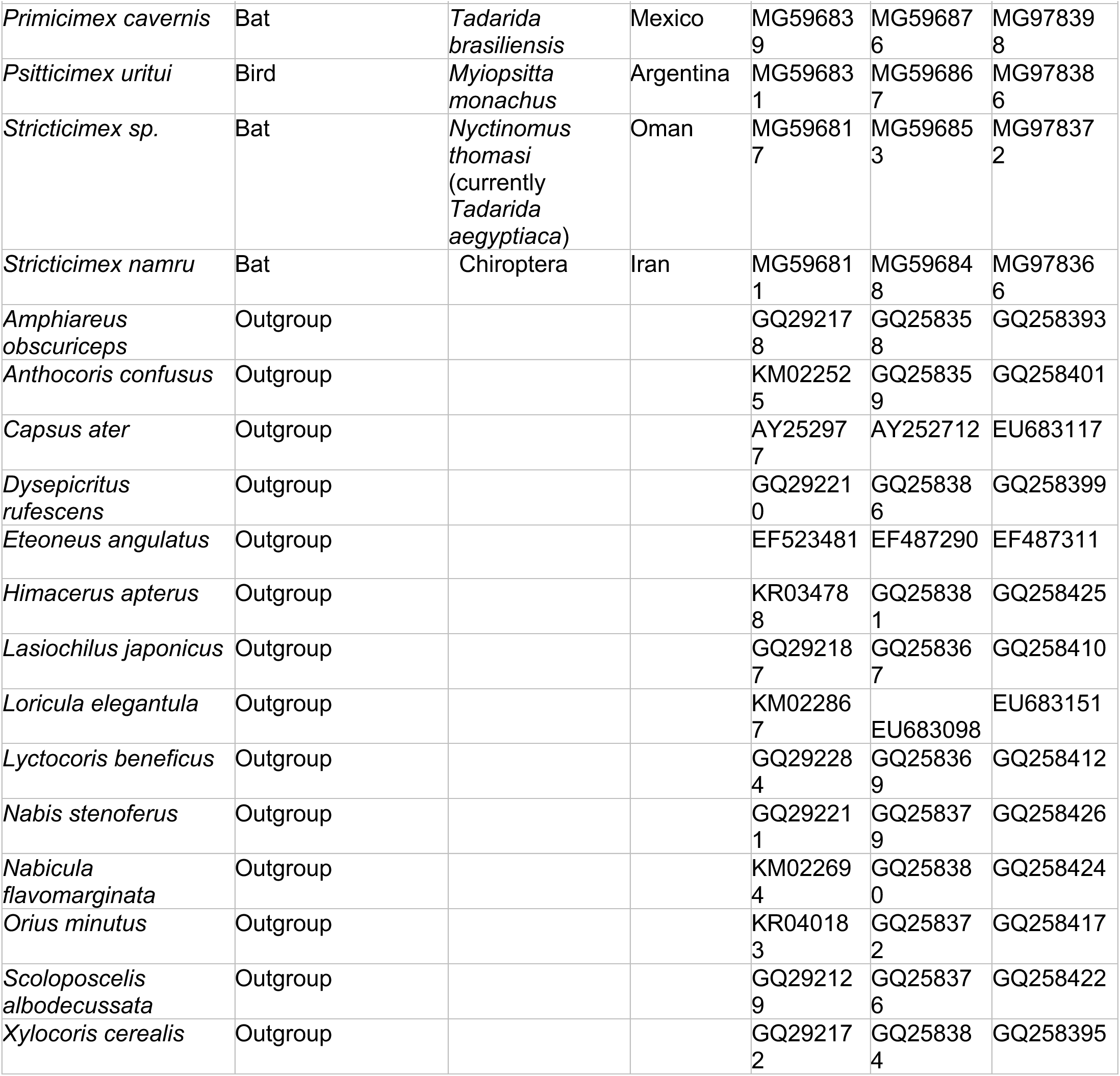
COI, 16S and 18S Sequences of Cimicidae species and outgroups obtained from NCBI GenBank.

| Species | Host group | Host species | Country | COI | 16S | 18S |
| --- | --- | --- | --- | --- | --- | --- |
| Cimicid 019 | Bat | *Mops plicatus* | Cambodia | PV273274 | PV275595 | PV275585 |
| Cimicid 049 | Bat | *Mops plicatus* | Cambodia | PV273269 | PV275596 | PV275586 |
| Cimicid 060 | Bat | *Mops plicatus* | Cambodia | PV273275 | PV275597 | PV275587 |
| Cimicid 070 | Bat | *Mops plicatus* | Cambodia | PV273276 | PV275598 | PV275588 |
| Cimicid 076 | Bat | *Mops plicatus* | Cambodia | PV273272 | PV275599 | PV275589 |
| Cimicid 077 | Bat | *Mops plicatus* | Cambodia | PV273273 | PV275600 | PV275590 |
| Cimicid 083 | Bat | *Mops plicatus* | Cambodia | PV273268 | PV275601 | PV275591 |
| Cimicid 084 | Bat | *Mops plicatus* | Cambodia | PV273270 | PV275602 | PV275592 |
| Cimicid 087 | Bat | *Mops plicatus* | Cambodia | PV273271 | PV275603 | PV275593 |
| *Acanthocrios furnarii* | Bird | *Furnarius rufus* (nest) | Brazil | MG596830 | MG596866 | MG978385 |
| *Afrochimex constrictus* | Bat | *Rousettus aegyptiacus* | Kenya | MG596805 | MG596842 | MG978358 |
| *Afrochimex constrictus* 2 | Bat | *Rousettus aegyptiacus* | Kenya | MG596806 | MG596843 | MG978359 |
| *Aphrania barys* | Bat | *Neoromicia capensis* (currently *Laephotis capensis*) | Namibia | MG596820 | MG596856 | MG978375 |
| *Aphrania barys* 2 | Bat | *Neoromicia capensis* (currently *Laephotis capensis*) | South Africa | MG596825 | MG596861 | MG978380 |
| *Aphrania elongata* | - | Unknown | Senegal | MG596812 | MG596849 | MG978367 |
| *Aphrania elongata* 2 | Bat | *Scotophilus leucogaster* | Mauritania | KF018763 | KF018729 | KF018715 |
| *Aphrania recta* | Bat | *Nycticeinops schlieffeni* | Mauritania | KF018764 | KF018730 | KF018716 |
| *Bucimex chilensis* | - | Unknown | Chile | MG596840 | MG596877 | MG978399 |
| Cacodminae gen. sp. | Bat | *Scotophilus leucogaster* | Mauritania | KF018764 | KF018730 | KF018716 |
| *Cacodmus sparsilis* | Bat | *Pipistrellus dhofarensis* | Oman | MG596813 | MG596850 | MG978369 |
| *Cacodmus vicinus* | Bat | *Scotoecus hirundo* | Senegal | MG596819 | MG596855 | MG978374 |
| *Cacodmus vicinus* 2 | Bat | *Pipistrellus* sp. | Spain | MG596816 | MG596852 | MG978371 |
| *Cacodmus villosus* | Bat | *Pipistrellus hesperidus* | Ethiopia | MG596821 | MG596857 | MG978376 |
| *Cacodmus villosus* 2 | Bat | *Neoromicia capensis* (currently *Laephotis capensis*) | Namibia | MG596823 | MG596859 | MG978378 |
| *Cimex adjunctus* | Bat | *Nycticeius humeralis* | USA | GU985536 | GU985558 | KF018712 |
| *Cimex emarginatus* | Bat | *Myotis alcathoe* cf. | Bulgaria | MG596837 | MG596874 | MG978396 |
| *Cimex emarginatus* 2 | Bat | *Pipistrellus pipistrellus* | Morocco | MF680526 | MF680517 | MG978397 |
| *Cimex hemipterus* | Human | *Homo sapiens* | Kenya | MG596826 | MG596862 | MG978381 |
| *Cimex hemipterus* 2 | Human | *Homo sapiens* | Malaysia | KF018754 | KF018724 | KF018710 |
| *Cimex latipennis* | Bat | *Myotis lucifugus* | Canada | KF018758 | KF018734 | KF018720 |
| *Cimex latipennis* 2 | Bat | *Myotis volans* | Canada | KF018757 | KF018733 | KF018719 |
| *Cimex lectularius* | Human | *Homo sapiens* | Czechia | GU985524 | GU985546 | KF018711 |
| *Cimex lectularius* 2 | Human | *Homo sapiens* | UK | MG596836 | MG596873 | MG978394 |
| *Cimex pipistrelli* | Bat | *Pipistrellus* sp. | UK | GU985534 | GU985556 | MG978393 |
| *Cyanolicimex patagonicus* | Bird | *Cyanoliseus patagonus* | Argentina | MG596833 | MG596869 | MG978388 |
| *Haematosiphon inodorus* | Bird | *Falco mexicanus* (nest) | USA | MG596829 | MG596865 | MG978384 |
| *Latrocimex spectans* | Bat | *Noctilio leporinus* | Belize | MW269881 | MW270938 | MZ378786 |
| *Leptocimex duplicatus* | - | Unknown | Israel | MG596810 | MG596847 | MG978365 |
| *Leptocimex inordinatus* | Bat | Chiroptera | Thailand | KT380161 | KT592538 | - |
| *Ornithocoris pallidus* | Bird | *Delichon urbicum* (nest) | USA | MG596828 | MG596864 | MG978383 |
| *Paracimex avium* | Bird | *Aerodramus salanganus* | Indonesia | MG596807 | MG596844 | MG978360 |
| *Paracimex chaeturus* | Bird | *Aerodramus brevirostris* | China | MF680531 | MF680520 | MG978362 |
| *Paracimex setosus* | Bird | *Aerodromus* sp. | Malaysia | KF018761 | KF018735 | KF018721 |
| *Primicimex cavernis* | Bat | *Tadarida brasiliensis* | Mexico | MG596839 | MG596876 | MG978398 |
| *Psitticimex uritui* | Bird | *Myiopsitta monachus* | Argentina | MG596831 | MG596867 | MG978386 |
| *Stricticimex sp.* | Bat | *Nyctinomus thomasi*<br>(currently *Tadarida aegyptiaca*) | Oman | MG596817 | MG596853 | MG978372 |
| *Stricticimex namru* | Bat | Chiroptera | Iran | MG596811 | MG596848 | MG978366 |
| *Amphiareus obscuriceps* | Outgroup |  |  | GQ292178 | GQ258358 | GQ258393 |
| *Anthocoris confusus* | Outgroup |  |  | KM022525 | GQ258359 | GQ258401 |
| *Capsus ater* | Outgroup |  |  | AY252977 | AY252712 | EU683117 |
| *Dysepicritus rufescens* | Outgroup |  |  | GQ292210 | GQ258386 | GQ258399 |
| *Eteoneus angulatus* | Outgroup |  |  | EF523481 | EF487290 | EF487311 |
| *Himacerus apterus* | Outgroup |  |  | KR034788 | GQ258381 | GQ258425 |
| *Lasiochilus japonicus* | Outgroup |  |  | GQ292187 | GQ258367 | GQ258410 |
| *Loricula elegantula* | Outgroup |  |  | KM022867 | EU683098 | EU683151 |
| *Lyctocoris beneficus* | Outgroup |  |  | GQ292284 | GQ258369 | GQ258412 |
| *Nabis stenoferus* | Outgroup |  |  | GQ292211 | GQ258379 | GQ258426 |
| *Nabicula flavomarginata* | Outgroup |  |  | KM022694 | GQ258380 | GQ258424 |
| *Orius minutus* | Outgroup |  |  | KR040183 | GQ258372 | GQ258417 |
| *Scoloposcelis albodecussata* | Outgroup |  |  | GQ292129 | GQ258376 | GQ258422 |
| *Xylocoris cerealis* | Outgroup |  |  | GQ292172 | GQ258384 | GQ258395 |

**Supplementary Table S2:** Body measurements of *Stricticimex phnomsampovensis* Suor & Maquart sp. Nov.

|  |  | Holotyp<br>e Male | Allotyp<br>e Female | Paratyp<br>e 1<br>Female | Paratyp<br>e 2<br>Male | Paratyp<br>e 3<br>Female | Paratyp<br>e 4<br>Female | Paratyp<br>e 5<br>Female<br>(juvenile<br>) |
| --- | --- | --- | --- | --- | --- | --- | --- | --- |
|  | Accession<br>n° | Ento_00<br>6 | Ento_00<br>1 | Ento_00<br>2 | Ento_00<br>6 | Ento_00<br>3 | Ento_00<br>4 | Ento_00<br>5 |
| **Head** | Lenght | 0,44 | 0,55 | 0,48 | 0,51 | 0,45 | 0,47 | 0,53 |
|  | Width | 0,41 | 0,51 | 0,49 | 0,48 | 0,53 | 0,54 | 0,45 |
|  | Eye | 0,1 | 0,09 | 0,08 | 0,76 | 0,067 | 0,07 | 0,07 |
|  | Interocular | 0,4 | 0,45 | 0,45 | 0,43 | 0,45 | 0,46 | 0,41 |
| **Antenna** | I | 0,15 | 0,15 | 0,15 | 0,16 | 0,13 | 0,14 | 0,15 |
|  | II | 0,42 | 0,45 | 0,46 | 0,47 | 0,47 | 0,45 | 0,43 |
|  | III | 0,7 | 0,83 | 0,79 | 0,78 | 0,67 | 0,75 | 0,58 |
|  | IV | 0,43 | 0,57 | 0,37 | 0,51 | 0,52 | 0,42 | 0,39 |
|  | **Total** | **1,7** | **2** | **1,77** | **1,92** | **1,79** | **1,76** | **1,55** |
| **Rostrum** | I | 0,2 | 0,23 | 0,2 | 0,2 | 0,2 | 0,2 | 0,17 |
|  | II | 0,18 | 0,21 | 0,17 | 0,2 | 0,2 | 0,2 | 0,16 |
|  | III | 0,1 | 0,13 | 0,1 | 0,13 | 0,1 | 0,17 | 0,08 |
|  | **Total** | **0,48** | **0,57** | **0,47** | **0,53** | **0,5** | **0,57** | **0,41** |
| **Pronotum** | Lenght | 0,28 | 0,3 | 0,27 | 0,3 | 0,28 | 0,3 | 0,23 |
|  | Width | 0,6 | 0,68 | 0,65 | 0,66 | 0,68 | 0,67 | 0,61 |
|  | Setae 1 | 0,16 | 0,17 | 0,17 | 0,17 | 0,19 | 0,17 | 0,17 |
|  | Setae 2 | 0,18 | 0,18 | 0,21 | 0,2 | 0,2 | 0,2 | 0,16 |
| **Hemelytra**<br>**s** | Lenght | 0,37 | 0,38 | 0,38 | 0,37 | 0,38 | 0,37 | Na |
|  | Width | 0,19 | 0,20 | 0,18 | 0,20 | 0,19 | 0,18 | Na |
| **Hind leg** | Femur | 0,79 | 0,94 | 0,83 | 0,81 | 0,88 | 0,93 | 0,75 |
|  | Tibia | 1,6 | 1,86 | 1,64 | 1,63 | 1,73 | 1,89 | 1,53 |
|  | Width F | 0,18 | 0,18 | 0,17 | 0,18 | 0,17 | 0,17 | 0,17 |

**Supplementary Table S3.** GenBank accession numbers.

| **RdRp Gene** |  |
| --- | --- |
| **Accession** | **Organism Name** |
| AAZ94069 | Aedes pseudoscutellaris reovirus |
| AAL31497 | Chum salmon reovirus CS |
| AAM92745 | Golden shiner reovirus |
| ADZ31977 | Scophthalmus maximus reovirus |
| ABV01040 | American grass carp reovirus |
| WPR16556 | Avian associated coltivirus |
| UPN67769 | California hare coltivirus |
| WAK76799 | Colorado tick fever virus |
| AAM18357 | Colorado tick fever virus |
| WHS68118 | Colorado tick fever virus |
| UVC65861 | Colorado tick fever virus |
| NP_690891 | Colorado tick fever virus |
| UVC65874 | Colorado tick fever virus |
| XCO66239 | Colorado tick fever virus |
| AAK73521 | Lymantria dispar cypovirus 1 |
| AAK73087 | Lymantria dispar cypovirus 14 |
| ACA53380 | Choristoneura occidentalis cypovirus 16 |
| AHJ14791 | Inachis io cypovirus 2 |
| AZG03007 | Eyach virus |
| AZG03006 | Eyach virus |
| AAM18359 | Eyach virus |
| AAM18358 | Eyach virus |
| ACF49423-ACF49425 | Eyach virus |
| AND77148 | Eyach virus |
| QYV43093 | Eyach virus |
| NP_620280 | Eyach virus |
| QKK82923 | Fennes virus |
| QIS88057 | Fennes virus |
| AAK40249 | Fiji disease virus |
| QYV43106 | Gierle tick virus |
| QZR93734 | Jeddah tick coltivirus |
| UYL95536 | Kashgar Reovi tick virus 1 |
| YP_010086008 | Kundal virus |
| WAK76791 | Lishui pangolin virus |
| WAK76790 | Lishui pangolin virus |
| QIE06438 | Lishui pangolin virus |
| URG17203 | Lishui pangolin virus P196T/pangolin/2018 |
| AMU04173 | Mahlapitsi orthoreovirus |
| ANG56321 | Maize rough dwarf virus |
| AAO73182 | Mal de Rio Cuarto virus |
| AAL36027 | Mammalian orthoreovirus 4 Ndelle |
| ANJ21365 | Mammalian orthoreovirus 1 |
| AGG40205 | Mammalian orthoreovirus 2 |
| ADJ00316 | Mammalian orthoreovirus 3 Dearing |
| AAP45577 | Cryphonectria parasitica mycoreovirus-1 (9B21) |
| BAC98431 | Mycoreovirus 3 |
| BFG88135 | Nakatsu tick virus |
| UYL95529 | Nanning Reovi tick virus 1 |
| UYL95538 | Nanning Reovi tick virus 2 |
| WHP37825 | Nelson Bay orthoreovirus |
| BAQ19494 | Pycnonotidae orthoreovirus |
| BAA08542 | Nilaparvata lugens reovirus |
| UJQ88099 | O'hara headland virus |
| UXL90843 | Qinghe tick reovirus |
| WWV88571 | Reoviridae sp. |
| WWV88567 | Reoviridae sp. |
| WWV88564 | Reoviridae sp. |
| WWV88562 | Reoviridae sp. |
| WWV88561 | Reoviridae sp. |
| WWV88558 | Reoviridae sp. |
| WAK76796 | Reoviridae sp. |
| WAK76797 | Reoviridae sp. |
| WWV87832 | Reoviridae sp. |
| WAK76800 | Reoviridae sp. |
| WAK76803 | Reoviridae sp. |
| WAK76785 | Reoviridae sp. |
| WAK76798 | Reoviridae sp. |
| WAK76806 | Reoviridae sp. |
| WWV88207 | Reovirus sp. |
| CAC82519 | Rice black streaked dwarf virus |
| AAC36456 | Rice ragged stunt virus |
| USH09528 | Salmon River virus |
| UVC65887 | Salmon River virus |
| QVL22803 | Shelly headland virus |
| UJQ88339 | Shelly headland virus |
| UJQ88340 | Shelly headland virus |
| UJQ88341 | Shelly headland virus |
| UJQ88342 | Shelly headland virus |
| UJQ88343 | Shelly headland virus |
| AYP67545 | Shelly headland virus |
| CBH31251 | Southern rice black-streaked dwarf virus |
| YP_010839657 | Tai Forest reovirus |
| BBA54722 | Tarumizu tick virus |
| YP_010086016 | Tarumizu tick virus |
| BBA54748 | Tarumizu tick virus |
| BBK20271 | Tarumizu tick virus |
| BDB06937 | Tarumizu tick virus |
| AOM63686 | chelonian orthoreovirus |
| UYL95532 | Yanbian Reovi tick virus 1 |
| UYL95533 | Yanbian Reovi tick virus 2 |
| UYL95534 | Yanbian Reovi tick virus 3 |
| CAG9164797 | Zeboroti virus |
| UYL95530 | Zhangjiakou Reovi tick virus 1 |
| WYW04511 | Reoviridae sp. |
| WYW04510 | Reoviridae sp. |
| WZL61382 | Calla Lily Valley virus |

| VP2 Gene |  |
| --- | --- |
| Accession | Organism Name |
| ACF49426 | Eyach virus |
| ACF49427 | Eyach virus |
| ACF49428 | Eyach virus |
| AND77149 | Eyach virus |
| AYP67546 | Shelly headland virus |
| BAS02070 | Cryphonectria parasitica mycoreovirus-1 (9B21) |
| BBA54723 | Tarumizu tick virus |
| BBA54749 | Tarumizu tick virus |
| BBK20272 | Tarumizu tick virus |
| BDB06936 | Tarumizu tick virus |
| BFG88136 | Nakatsu tick virus |
| NP_620281 | Eyach virus |
| NP_690892 | Colorado tick fever virus |
| QIE06439 | Lishui pangolin virus |
| QIS88058 | Fennes virus |
| QYV43094 | Eyach virus |
| QYV43107 | Gierle tick virus |
| QZR93735 | Jeddah tick coltivirus |
| UJQ88344 | Shelly headland virus |
| UPN67770 | California hare coltivirus |
| USH09529 | Salmon River virus |
| UVC65862 | Colorado tick fever virus |
| UVC65875 | Colorado tick fever virus |
| UVC65888 | Salmon River virus |
| UXL90844 | Qinghe tick reovirus |
| WHS68119 | Colorado tick fever virus |
| WKE34862 | Colorado tick fever virus |
| WKE34869 | Colorado tick fever virus |
| WKE34874 | Colorado tick fever virus |
| WKE34884 | Colorado tick fever virus |
| WKE34901 | Colorado tick fever virus |
| WKF25357 | Colorado tick fever virus |
| WKF25359 | Colorado tick fever virus |
| WWV87811 | Reoviridae sp. |
| WWV87834 | Reoviridae sp. |
| WWV88557 | Reoviridae sp. |
| WWV88560 | Reoviridae sp. |
| WWV89314 | Riboviria sp. |
| WZL61383.1 | Calla Lily Valley virus |
| XCO66240.1 | Colorado tick fever virus |
| XKB76581 | Reovirales sp. |
| XKB76582 | Reovirales sp. |
| YP_001936005 | Mycoreovirus 1 |
| YP_009252406 | Sclerotinia sclerotiorum mycoreovirus 4 |
| YP_010086009 | Kundal virus |
| YP_010086015 | Tarumizu tick virus |
| YP_010839655 | Tai Forest reovirus |
| YP_392476 | Mycoreovirus 3 |
| NC_007154.1 | Fiji disease virus |

